# A cohort of *Caenorhabditis* species lacking the highly conserved *let-7* microRNA

**DOI:** 10.1101/2020.11.10.377101

**Authors:** Charles Nelson, Victor Ambros

## Abstract

*let-7* is a highly conserved microRNA with critical functions integral to cell fate specification and developmental progression in diverse animals. In *Caenorhabditis elegans, let-7* is a component of the heterochronic (developmental timing) gene regulatory network, and loss-of-function mutations of *let-7* result in lethality during the larval to adult transition due to misregulation of the conserved *let-7* target, *lin-41*. To date, no bilaterian animal lacking *let-7* has been characterized. In this study, we identify a cohort of nematode species within the genus *Caenorhabditis*, closely related to *C. elegans*, that lack the *let-7* microRNA, owing to absence of the *let-7* gene. Using *C. sulstoni* as a representative *let-7*-lacking species to characterize normal larval development in the absence of *let-7*, we demonstrate that, except for the lack of *let-7*, the heterochronic gene network is otherwise functionally conserved. We also report that species lacking *let-7* contain a group of divergent *let-7* orthologs -- also known as the *let-7-family* of microRNAs -- that have apparently assumed the role of targeting the *lin-41* mRNA.

**Summary Statement:** We have identified a group of *Caenorhabditis* species that lack *let-7a*, an otherwise highly conserved and nearly ubiquitous microRNA that was previously thought to be critical to bilaterian animal development.

## INTRODUCTION

MicroRNAs are approximately 22 nucleotide (nt) non-coding RNAs that negatively regulate protein expression through base pairing of nucleotides 2-8 of the microRNA (known as the microRNA seed) to complementary sequences in target mRNA 3’ UTRs. Base pairing with nucleotides 9-22 can also contribute to target repression but non-seed base pairing is less constrained than seed pairing. Accordingly, evolutionary conservation of microRNA sequences is generally highest for nucleotides 2-8, and less so for non-seed nucleotides (Bartel, 2009; Ambros and Ruvkun, 2018).

Unlike most microRNAs, the entirety of nucleotides 1-22 of *let-7* are highly conserved across bilaterians (Fig. 1A) (Pasquinelli et al., 2000). Why *let-7* non-seed sequences are so deeply conserved remains a mystery. Alongside the deep conservation of the entire *let-7* sequence, *let-7* microRNA function is also conserved; across diverse animal phyla, *let-7* expression coincides with differentiation and opposes stem cell pluripotency (Reinhart et al., 2000; Pasquinelli et al., 2000; Roush and Slack, 2008; Balzeau et al., 2017). Accordingly, in certain contexts *let-7* functions as a tumor suppressor through restricting the expression of proteins involved in cell proliferation, growth, and metabolism, among others (Balzeau et al., 2017). We believe that the deep conservation of *let-7* sequence holds secrets to important evolutionarily conserved molecular interactions vital to *let-7* function, and therefore, a better understanding of *let-7* conservation will reveal insights into mechanisms of microRNA function and regulation.

**Figure 1.**
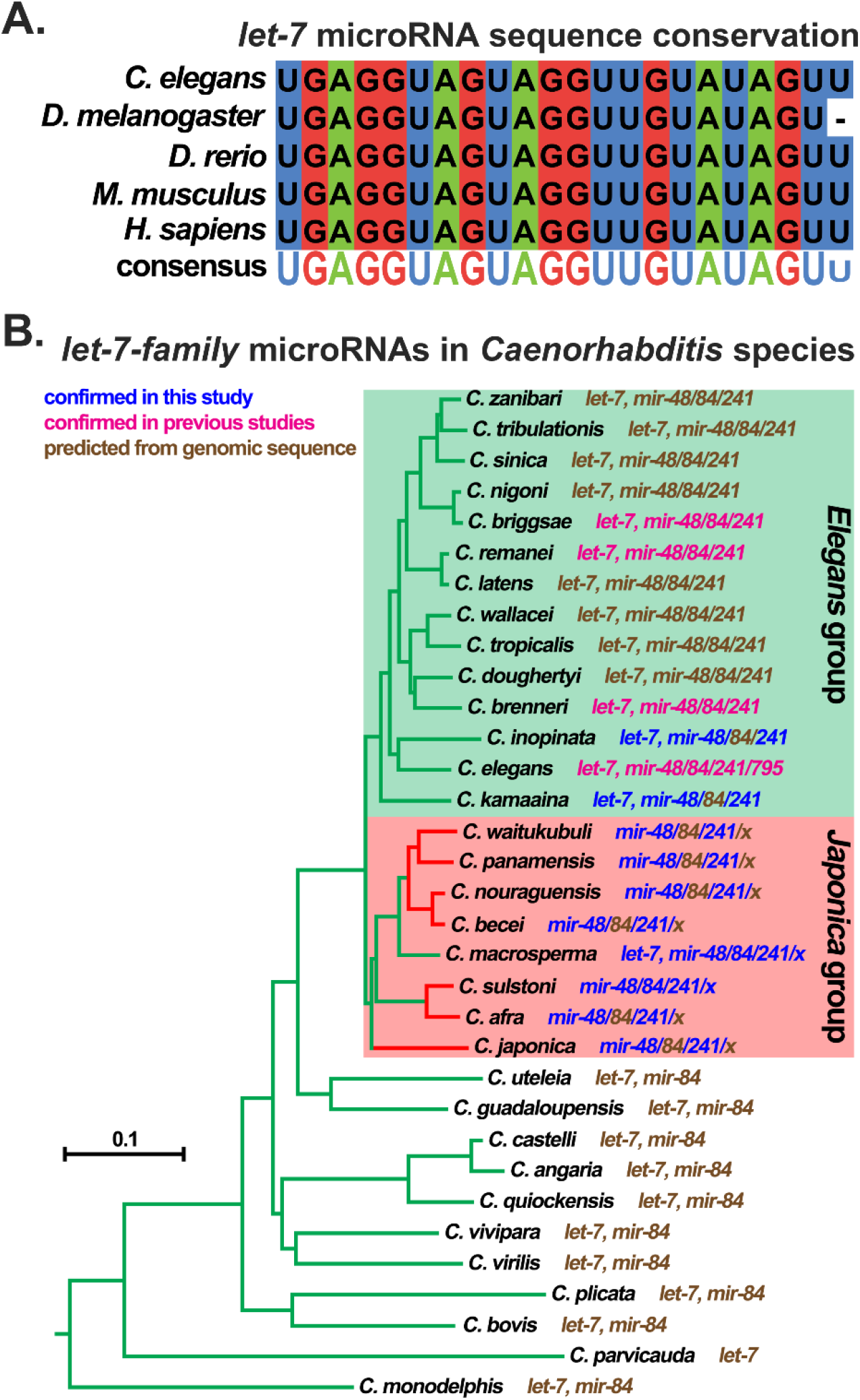
Conservation of the *let-7* microRNA. (A) Sequence conservation of *let-7* microRNA. (B) Phylogenetic tree of *Caenorhabditis* species based on phylogeny from Stevens et al., 2020. For each species, the makeup of their *let-7-family* microRNAs is indicated. *let-7-family* microRNAs are predicted from their genomic sequences (brown font), detection in previous studies (magenta font) (Pasquinelli et al., 2000; Reinhart et al., 2000; Lau et al., 2001; Lim et al., 2003; Ruby et al., 2006; de Wit et al., 2009; Shi et al., 2013), or detection in this study (blue font). Green branches highlight species that are either predicted or confirmed to have *let-7.* Red branches highlight species that are either predicted or confirmed to lack *let-7*. Outlined in the green box are species recovered in the *Elegans* group. Outlined in the red box are species recovered in the *Japonica* group. Branch lengths are in substitutions per site. Scale bar is shown.

In most bilaterian animal clades (i.e. all animals with bilaterian symmetry), including mammals and nematodes, the *let-7* gene family has been expanded into a number of related microRNAs that share the same seed sequence but differ in their non-seed nucleotides (Roush and Slack, 2008). In *C. elegans*, the *let-7-family* consists of *let-7, mir-48, mir-84, mir-241*, and *mir-795* (Abbott et al., 2005). Despite the presence of multiple *let-7-family* microRNAs (which could in principle substitute for one another by seed-based pairing to targets), most animals have nevertheless retained the original *let-7* microRNA (specifically, *let-7a*) with its non-seed nucleotide sequence conserved (Pasquinelli et al., 2000; Roush and Slack, 2008).

The *C. elegans* heterochronic gene (developmental timing) regulatory network consists of genes that either promote or restrict developmental cell fate progression. Integrated into the heterochronic pathway are the protein coding genes *lin-14, lin-28, lin-29, lin-41, lin-46*, and *hbl-1*, as well as the microRNAs *lin-4* and the *let-7-family* (*let-7, mir-48, mir-84*, and *mir-241*). Loss-of-function mutations of genes that restrict cell fate progression, such as *lin-14, lin-28, lin-41*, or *hbl-1*, results in precocious development of the hypodermis through the skipping of larval-specific cellular events; whilst loss-of-function mutations of genes that promote cell fate progression, such as *lin-29, lin-46, lin-4*, or the *let-7-family*, results in retarded development of the hypodermis through the reiteration of larval-specific cellular divisions (Chalfie et al., 1981; Ambros and Horvitz, 1984; Ambros, 1989; Fay et al., 1999; Reinhart et al., 2000; Slack et al., 2000; Lin et al., 2003; Pepper et al., 2004; Abbott et al., 2005).

In *C. elegans*, expression of the *lin-4* microRNA increases during the L1 and L2 stages to promote the transition from L1 cell fates to later larval cell fates through its targeted repression of synthesis of the LIN-14 and LIN-28 proteins (which promote L1 and L2 larval stage cell fates) by base-pairing to the 3’ UTRs of *lin-14* and *lin-28* mRNAs (Chalfie et al., 1981; Ambros and Horvitz, 1984; Ambros, 1989; Lee et al., 1993; Moss et al., 1997; Lim et al., 2003). Similarly, *mir-48, mir-84*, and *mir-241* (*mir-48/84/241*), whose expressions increase during the L2 and L3 larval stages to promote transitions to later larval stages by negatively regulating *lin-14, lin-28*, and *hbl-1* through base-pairing to their respective 3’ UTRs (Abrahante et al., 2003; Lin et al., 2003; Abbott et al., 2005; Tsialikas et al., 2017).

In *C. elegans, let-7* is required for cell cycle exit and differentiation in hypodermal cell lineages at the end of larval development, which is reflected by the dramatic up regulation of mature *let-7* microRNA in terminal larval stages (Reinhart et al., 2000). *let-7* function and regulation are deeply integrated into the *C. elegans* heterochronic gene regulatory network; LIN-28 post transcriptionally restricts *let-7* microRNA biogenesis to later larval stages, and *let-7* negatively regulates the evolutionarily conserved pluripotency-promoting genes *lin-41* and *lin-28* through base-pairing to the 3’ UTRs of the *lin-41* and *lin-28* mRNAs (Reinhart et al., 2000; Slack et al., 2000; Vella et al., 2004; Ding and Grosshans, 2009; Lehrbach et al., 2009; Van Wynsberghe et al., 2011; Ecsedi et al., 2015; Stefani et al., 2015). The reciprocal direct regulation between *let-7* and LIN-28, and the direct regulation of *lin-41* by *let-7*, are evolutionarily conserved across vertebrates and invertebrates, suggesting that the unusual conservation of *let-7* sequence may be related to these conserved intimate interactions of *let-7* microRNA with the *lin-28* and *lin-41* mRNAs and with LIN-28 protein (Reinhart et al., 2000; Slack et al., 2000; Kloosterman et al., 2004; Vella et al., 2004; Schulman et al., 2005; Lin et al., 2007; Heo et al., 2008; Newman et al., 2008; Rybak et al., 2008; Viswanathan et al., 2008; Lehrbach et al., 2009; Nam et al., 2011; Van Wynsberghe et al., 2011; Piskounova et al., 2011; Stratoulias et al., 2014; Ecsedi et al., 2015; Stefani et al., 2015; Balzeau et al., 2017).

Despite the sharing of a common seed sequence and the seemingly redundant potential to regulate the same targets, the *let-7-family* microRNAs (*mir-48/84/241*) and *let-7* do not function interchangeably. Specifically, *mir-48/84/241*, as a semi-redundant cohort, primarily regulate *lin-14, lin-28*, and *hbl-1* during L1-L3 cell fate transitions, whereas *let-7* regulates *lin-41* during a larval to adult cell fate switch (Slack et al., 2000; Vella et al., 2004; Abbott et al., 2005; Ecsedi et al., 2015; Tsialikas et al., 2017; Ilbay and Ambros, 2019). *let-7* loss-of-function (*let-7(lf)*) mutants display phenotypes distinguishable from those of triply-mutant *mir-84(lf)*; *mir-48(lf)mir-241(lf)* animals, as a consequence of the stage-specific de-repression of their respective targets – essentially gain-of-function of *lin-41*, or gain-of-function *lin-14/lin-28/hbl-1*, respectively (Abbott et al., 2005; Aeschimann et al., 2019). The specificity of *let-7* for regulation of *lin-41* is thought to be conferred by base pairing of 3’ non-seed sequences of *let-7* to the *let-7* complementary sites in the *lin-41* 3’ UTR (Reinhart et al., 2000; Vella et al., 2004; Ecsedi et al., 2015), suggesting that conservation of *let-7* non-seed sequences could reflect evolutionary pressure to conserve functional distinctions between *let-7* and other *let-7-family* members.

The deep evolutionary roots of *let-7* in the heterochronic pathway, including the apparent conservation of specific targeting of *lin-41* by *let-7* suggests that hypothetical evolutionary loss of *let-7* could be expected to be accompanied by significant divergence, compared to *C. elegans*, in the functions of heterochronic genes downstream of *let-7*, such as *lin-41* and *lin-29*, and/or upstream genes, such as *lin-14, lin-28*, and *hbl-1.* Exploration of these questions would require the identification of species closely related to *C. elegans* that lack *let-7.*

Here we identify a faction of *Caenorhabditis* species within the *Japonica* group, a sister group to the *Elegans* group, that lack *let-7*. As far as we know, this is the first described instance of two sister clades where all known species of one clade have retained *let-7*, while numerous species of the sister clade do not have *let-7.* We demonstrate that for an exemplary *let-7*-lacking species, *C. sulstoni*, the functional architecture of the heterochronic pathway is otherwise conserved compared to *C. elegans.* Our findings indicate that *lin-41* is apparently regulated by the remaining *let-7-family* microRNAs in most *Japonica* group species, suggesting that the heterochronic pathway can evolve to re-delegate *let-7-family* function under certain evolutionary circumstances.

## RESULTS

### Most *Caenorhabditis* species belonging to the *Japonica* group lack *let-7* microRNA

The *let-7* microRNA was the first microRNA whose sequence and developmental function in promoting the differentiation of cell fates were shown to be conserved from nematodes to vertebrates (Fig. 1A) (Pasquinelli et al., 2000; Reinhart et al., 2000; Lin et al., 2007; Caygill and Johnston, 2008; Sokol et al., 2008). In the course of studying the evolution of regulatory sequences within the *let-7* locus of related nematodes, we discovered that most species of the *Japonica* group of *Caenorhabditis* lack the *let-7* sequence in their genomic assemblies (Fig. 1B; Table S1). Interestingly, one exceptional *Japonica* group species, *C. macrosperma*, had *let-7* sequence in its genomic assembly, which we confirmed using PCR amplification and Sanger sequencing (Fig. S1).

To determine if the lack of *let-7* sequence in the genomic assemblies of these *Japonica* group species could reflect major genomic rearrangements and/or anomalous assembly of genomic sequence, we analyzed the genome sequences surrounding *let-7* in *C. elegans* for potential synteny to corresponding genomic sequences of all *Caenorhabditis* species predicted to lack the *let-7* sequence. We found that the genomic assemblies of all the *Caenorhabditis* species lacking *let-7* sequence contain a region syntenic to the region surrounding the *let-7* locus of *C. elegans* (representative synteny shown in Fig. 2A). With the exception of *C. afra*, none of the *Caenorhabditis* species lacking *let-7* sequence exhibited any indication that loss of *let-7* sequence is associated with genomic rearrangement (Fig. S2).

**Figure 2.**
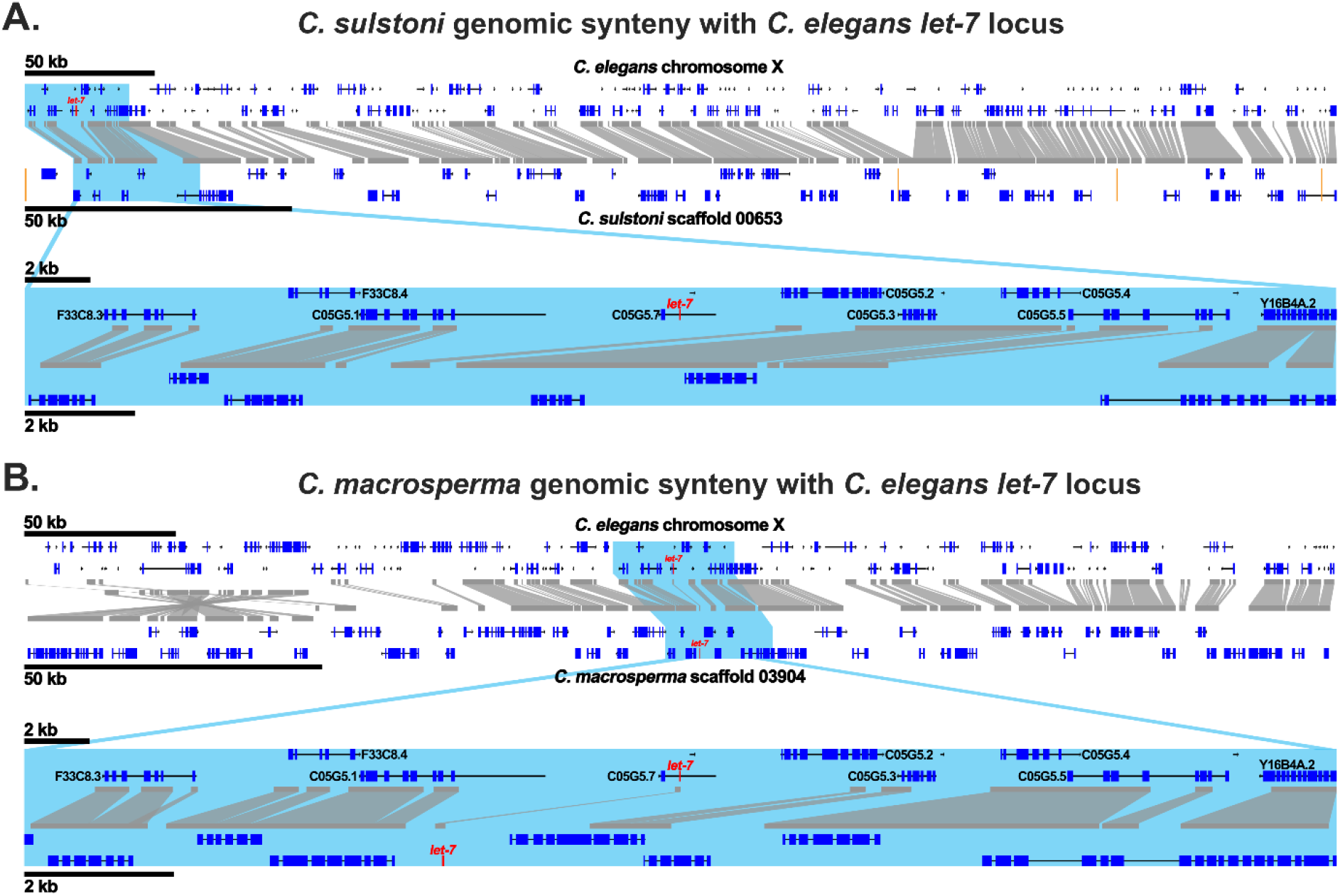
The genomic region containing the *let-7* sequence in *C. elegans* is syntenic to genomic regions in *C. sulstoni* and *C. macrosperma*. Synteny of a portion of *C. elegans* chromosome X containing the *let-7* sequence with *C. sulstoni* scaffold 00653 (A) and with *C. macrosperma* scaffold 03904 (B). Regions with sequence similarity are outlined in gray. Sequence gaps are shown in orange. *let-7* is shown in red. Highlighted in light blue are zoomed in views of the *C. elegans let-7* locus and its respective syntenies with *C. sulstoni* (A) and *C. macrosperma* (B).

The absence of the *let-7* sequence in the genomic assemblies of *Japonica* group species could reflect incomplete sequence coverage. In such cases, *let-7* could be expressed from DNA that was, for some reason, not detected by genomic sequencing. Therefore, to gather evidence, independent of genomic sequence, for whether or not *let-7* is expressed in *Japonica* group species, we profiled microRNAs using FirePlex miRNA assays in RNA samples from mixed-stage populations of eight *Japonica* species available from the *Caenorhabditis* Genetics Center (CGC), *C. waitukubuli, C. panamensis, C. nouraguensis, C. becei, C. macrosperma, C. sulstoni, C. afra*, and *C. japonic*a.

For all seven of the *Japonica* group species that lack *let-7* in their genome assemblies as well as *C. macrosperma*, which has *let-7* sequence in its genome assembly (Fig. 1B, 2B and S1; Table S1), we failed to detect *let-7* expression by FirePlex assay. By contrast, *let-7* was readily detectable using FirePlex in mixed-staged total RNA samples from species of the closely related *Elegans* group, *C. inopinata, C. elegans*, and *C. kamaaina* (Fig. 1B; Table S1). These FirePlex data also confirmed the expression of other microRNAs, including *lin-4* and the *let-7-family* microRNAs *mir-48* and *mir-241* in all eight *Japonica* group species, as well as in the three *Elegans* group species (Fig. 1B; Table S1).

### *C. macrosperma* expresses *let-7*

*C. macrosperma* is the one exceptional *Japonica* species in our experimental set that does contain *let-7* genomic sequence. Despite confirming the presence of the *let-7* sequence in the *C. macrosperma* genomic sequence assembly and its synteny to the *C. elegans let-7* genomic locus (Fig. S1 and Fig. 2B), we failed to detect *let-7* expression using the FirePlex assay. The FirePlex assay employs a panel of hybridization probes complementary to *C. elegans* microRNAs, which allows detection of microRNAs that precisely match the corresponding probe, or in some cases, that differ by a single internal nucleotide. While the *C. macrosperma* genomic sequence assembly contains an apparent *let-7-5p* guide RNA sequence identical to the *C. elegans let-7* microRNA, it is possible that *let-7* was not detected by FirePlex in our *C. macrosperma* RNA samples owing to significant 5’ or 3’ end variation and/or expression levels below the limits of detection by FirePlex. Similarly, other microRNAs that were not detected in our FirePlex data (such as the *let-7* paralog *mir-84*, which was detected in *C. elegans* only) (Fig. 1B; Table S1), could have been missed owing to low expression levels and/or significant interspecific variation in their non-seed sequences.

To definitively determine if *let-7* is expressed in *C. macrosperma*, we performed small RNA sequencing of RNA samples from each larval stage as well as early adult stage for both *C. elegans* and *C. macrosperma* (staging times shown in Fig. S3A and S3B). From these data, we confirmed that *let-7* is indeed expressed in *C. macrosperma* (Fig. 1B; Table S1). Unlike *C. elegans let-7*, whose expression dramatically increases during the L3 stage and peaks at the L4 stage and accumulates to levels similar to *mir-48* and *mir-84, C. macrosperma let-7* undergoes a more gradual and blunted increase in level during *C. macrosperma* development and never exceeds the level of any other *let-7-family* microRNA (Fig. 3A and 3B).

**Figure 3.**
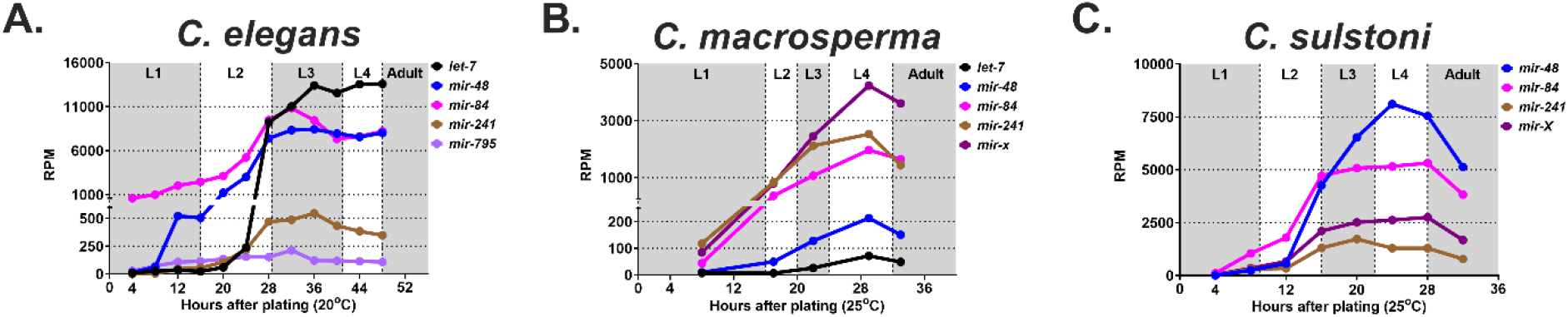
*let-7-family* microRNA temporal expression patterns for *C. elegans, C. macrosperma*, and *C. sulstoni*. Small RNA sequencing data showing expression of *let-7-family* microRNAs throughout *C. elegans* (A), *C. macrosperma* (B), and *C. sulstoni* (C) development. RPM refers to reads per million.

Small RNA sequencing also confirmed the expression of *C. macrosperma lin-4*, as well as the *let-7* family microRNAs, *mir-48, mir-241, mir-84* (which is one nucleotide shorter and has four internal nucleotide differences from *C. elegans mir-84*), and a novel *let-7-family* microRNA, *mir-X* (Fig. 1B, 3B and S4B; Table S1). Interestingly, the *C. macrosperma mir-X* genomic sequence is located 106 bp downstream of the *mir-84* sequence, suggesting that *mir-84* and *mir-X* may be produced from a common primary transcript. Similar to the *C. elegans let-7* family microRNAs, the developmental expression profiles of *mir-48/84/241/X* in *C. macrosperma* consist of gradual increases during the L1 to L2 stages and peaks during the L3 and L4 stages (Fig 3B). We note a relatively low expression of *mir-48* in *C. macrosperma* compared to *C. elegans mir-48* (Fig. 3A and 3B).

The temporal expression profile of *C. macrosperma lin-4* was similar to that of *C. elegans*, consisting of an increase in expression during early larval stages, followed by a broad peak during middle stages, and a decrease around the L4/adult transition (Fig. S4A and S4B).

### Robust assignment of *C. macrosperma* to the *Japonica* group

*C. macrosperma* was the only member of the *Japonica* group analyzed here to contain a *let-7* gene in its genome, implying the presence of *let-7* in a *Japonica* ancestor and loss of *let-7* during the divergence of these *Japonica* species. An alternative explanation for the exceptionalism of *C. macrosperma* as a *let-7*-containing *Japonica* species could be an erroneous phylogenic assignment of *C. macrosperma.* In this regard, *C. macrosperma* was first recovered as a member the *Japonica* group based on a comparison of selected gene segments across *Caenorhabditis* species (Kiontke et al., 2011; Felix et al., 2014). Subsequent genome-wide analyses that include additional *Caenorhabditis* species again recovered *C. macrosperma* in the *Japonica* group with maximal support (a Bayesian posterior probably of 1.0 and a bootstrap value of 100, respectively) (Stevens et al., 2019; Stevens et al., 2020), indicating a high likelihood of its correct assignment.

To obtain additional evidence confirming the evolutionary affinity of *C. macrosperma* with the *Japonica* group, we examined the phylogeny of the conserved *let-7* target gene, *lin-41*. We reasoned that if the presence of a *let-7* gene reflects evolutionary affinity of *C. macrosperma* with a clade outside *Japonica*, then such hypothetical divergence from *Japonica* might be reflected in the gene-specific phylogeny of *lin-41*, a conserved *let-7*-specific target. When we constructed a *lin-41* gene tree across 33 *Caenorhabditis* species, we observed a *lin-41* phylogeny nearly identical to the species phylogeny (Fig. S5), confirming the *Japonica* affinity of *C. macrosperma lin-41*, and strongly supporting the conclusion that *C. macrosperma* is an exceptional *Japonica* group species that has retained *let-7*.

Further support for the assignment of *C. macrosperma* to the *Japonica* group comes from considering the novel *let-7-family* member, *mir-x*, which we initially identified in small RNA sequencing from *C. macrosperma.* Nucleotide BLAST search analyses of *Caenorhabditis* species genomes revealed *mir-x* homologs to be present in all eight *Japonica* group species and in no species outside of the *Japonica* group (Fig. 1B; Table S1). This suggests that *mir-x* is a *Japonica* group specific *let-7-family* microRNA, and its presence in *C. macrosperma* supports the view that *C. macrosperma* is indeed a member of the *Japonica* group.

### The *mir-84* loci in the *Japonica* group are polycistronic with novel *let-7-family* microRNAs

The identification of *mir-x* within the *mir-84* locus of *C. macrosperma* led us to explore if there are more predicted novel *let-7-family* microRNAs in the *mir-84* loci of other *Caenorhabditis* species. To do this, we searched the genomic sequence surrounding the *mir-84* sequence for the presence of the *let-7-family* seed sequence (GAGGUAG) in all *Caenorhabditis* species. We next used *in silico* RNA folding to predict if the RNA would fold into a stem-loop structure indicative of a microRNA precursor. From this analysis, we concluded that all members of the *Japonica* group have between one and five extra *let-7-family* microRNAs in their *mir-84* loci. Interestingly, *mir-84* is always the 5’ most *let-7-family* microRNA in these loci, and these additional *let-7-family* microRNAs are most likely polycistronic with *mir-84* as they are no further than 221 nt apart (average of 116 ± 40 nt). We found no evidence of additional *let-7-family* microRNAs in the *mir-84* loci in any *Caenorhabditis* species outside of the *Japonica* group (Fig. 1B; Table S1).

### *C. sulstoni* serves as a representative species lacking *let-7*

*C. sulstoni* is a gonochoristic bacteriovore that was isolated from the feces of the east African millipede *Archispirostreptus gigas* purchased at an insect market in Berlin in spring 2013 (Stevens et al., 2019). Based on its robust and consistent developmental trajectory and ease of experimental manipulation (including sensitivity to RNAi), we adopted *C. sulstoni* as a representative *let-7*-lacking species for our experiments. *C. sulstoni* grows on standard nematode growth media (NGM) plates seeded with *E. coli*. *C. sulstoni* larval development appears largely similar to *C. elegans*, containing six distinct phases: an embryonic stage, L1 through L4 larval stages, and adult stage. The overall organismal morphology and cellular anatomy of *C. sulstoni* males and females appear markedly similar to the corresponding sexes of *C. elegans.* Particularly, the development of the *C. sulstoni* hypodermal cell lineages, the morphologies of stage specific cuticles, and the ontogeny of the gonad and vulva are similar to *C. elegans.*

Like *C. elegans, C. sulstoni* embryos can be isolated by sodium hypochlorite/sodium hydroxide treatment of populations that include gravid adults. After allowing embryos to hatch in the absence of food, populations of developmentally arrested L1 larvae (L1 diapause) are obtained. Addition of food to a population of L1 diapause larvae triggers synchronous initiation of larval development, thereby enabling the preparation of populations of developing larvae of defined larval stages for biochemical and molecular experiments. Unlike *C. elegans*, which is typically cultured between 15°C and 25°C, *C. sulstoni* can be cultured between 20°C and 30°C (Fig. S3A and S3C). Interestingly, at 25°C *C. sulstoni* develop from the L1 to adulthood 13 hours faster than *C. elegans*, 28 hours versus 41 hours, respectively (Fig. S3A and S3C). Moreover, at 30°C and 33°C (temperatures that do not support *C. elegans* development), *C. sulstoni* develops from the L1 stage to adulthood in approximately 23 hours (Fig. S3A and S3C). 15°C and 33°C seem to define the low and high limits of *C. sulstoni* temperature tolerance under our culture conditions; *C. sulstoni* can develop from L1 to adulthood at 15°C or 33°C but are sterile when raised at either temperature. Populations of *C. sulstoni* grown at 15°C become asynchronous and take approximately six days for the first animals to reach adulthood (Fig. S3C).

### Except for the lack of *let-7*, the heterochronic pathway is functionally conserved in *C. sulstoni*

In *C. elegans, let-7* functions as a significant component of the heterochronic pathway by ensuring proper developmental cell fate progression, particularly during the larval to adult cell fate transition (Fig. S6A) (Reinhart et al., 2000). Because this hallmark microRNA of the heterochronic pathway is lacking in *C. sulstoni*, we sought to determine the status of other major components of the heterochronic pathway. As mentioned in the introduction, the protein coding genes *lin-14, lin-28, lin-46, hbl-1, lin-41*, and *lin-29* as well as the microRNAs *lin-4, mir-48/84/241*, and *let-7* are critical components of the heterochronic pathway in *C. elegans* (Fig. S6A). In *C. elegans*, loss of *lin-14, lin-28, hbl-1*, or *lin-41* results in precocious development and the subsequent early formation of adult-specific structures including adult lateral alae (Ambros and Horvitz, 1984; Fay et al., 1999; Slack et al., 2000; Abrahante et al., 2003; Lin et al., 2003). By contrast, loss of *lin-4, lin-46, mir-48/84/241*, or *lin-29* result in retarded development, characterized by incomplete formation of adult-specific structures including adult lateral alae (Chalfie et al., 1981; Ambros, 1989; Ambros and Horvitz, 1984; Pepper et al., 2004; Abbott et al., 2005).

To assess the conservation of heterochronic pathway gene function between *C. elegans* and *C. sulstoni*, we first characterized the larva-to-adult cell fate transition in the hypodermis of *C. sulstoni.* During the L4-adult transition in *C. elegans*, hypodermal seam cells, which consist of a longitudinal string of cells of either side of the animal, finish dividing, fuse with each other to form a lateral line syncytium, and produce the adult-specific cuticular structure called lateral alae, which signifies the terminal differentiation of the seam cells (Sulston and Horvitz, 1977; Ambros and Horvitz, 1984). Similar to *C. elegans*, in *C. sulstoni* we observed seam cell fusion and the formation for adult alae during the L4-adult transition (Fig. 4A–4D).

**Figure 4.**
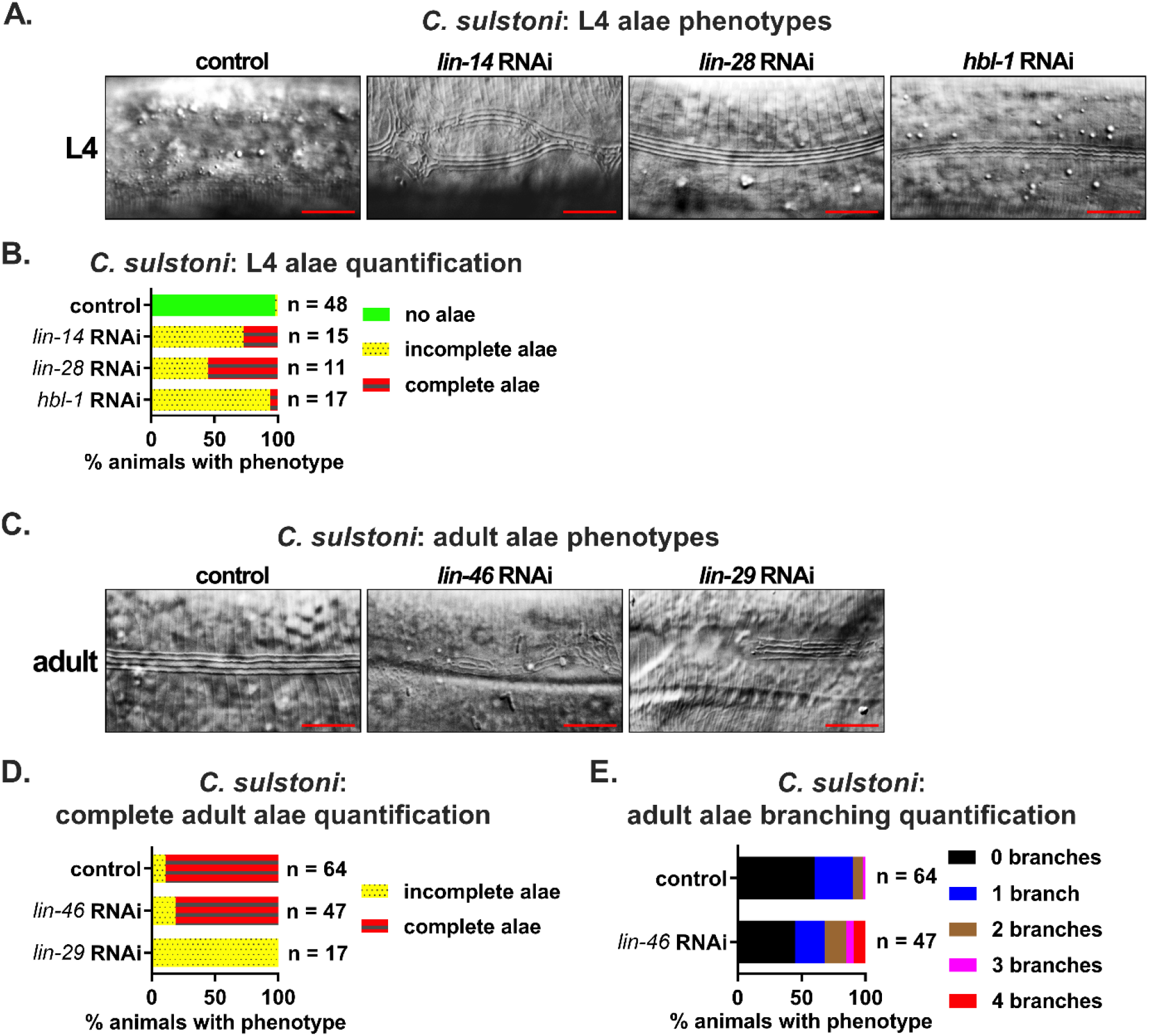
Heterochronic phenotypes associated with RNAi of *C. sulstoni lin-14, lin-28, hbl-1, lin-46*, and *lin-29*. (A and B) Panels from left to right: representative DIC images of *C. sulstoni* L4 hypodermis of animals fed control (empty vector), *lin-14, lin-28*, and *hbl-1* RNAi, respectively (A), and quantification of observed L4 alae phenotypes (B). Scale bars = 10 μM. (C-E) Panels from left to right: representative DIC images of *C. sulstoni* adult hypodermis of animals fed control (empty vector), *lin-46*, and *lin-29* RNAi, respectively (C), and quantification of observed complete adult alae (D) and adult alae branching (E) phenotypes observed. Scale bars = 10 μM. Note: the control (empty vector) and *lin-46* RNAi animals used for panel D were also used for panel E.

From our FirePlex microRNA profiling, we had already confirmed the expression of *lin-4* and the *let-7-family* microRNA genes *mir-48* and *mir-241* in *C. sulstoni* (Fig. 1B; Table S1). We sought to determine if these microRNAs are integrated into the *C. sulstoni* heterochronic network through their targeted repression of the *lin-14, lin-28*, and *hbl-1* 3’UTRs as they are in *C. elegans.* Because 3’ UTRs are not annotated in *C. sulstoni*, we used the *C. elegans* 3’ UTRs as a framework and examined sequences downstream of each respective gene’s stop codon for *lin-4* and *let-7-family* complementary sites (Table S2). For the *C. sulstoni lin-14, lin-28*, and *hbl-1* 3’ UTR regions, predicted sites for *lin-4* and *let-7-family* microRNAs were identified, indicating conservation of the targeting of these heterochronic genes by *lin-4* and *let-7-family* between *C. elegans* and *C. sulstoni* (Table S2; Fig. S6B).

As mentioned previously, the FirePlex assay we used to detect *lin-4* and *mir-48/241* in *C. sulstoni* employs a panel of hybridization probes complementary to *C. elegans* microRNAs, which does not allow for the detection of microRNAs with multiple nucleotide differences. This would include *C. sulstoni mir-84*, which, based on homology to *C. macrosperma*, is predicted to be one nucleotide shorter and have three internal nucleotide differences compared to *C. elegans mir-84*.

To assay for *mir-84* and to potentially identify novel *let-7-family* members in *C. sulstoni*, and to determine their temporal expression patterns during development, we performed small RNA sequencing of RNA samples from each larval stage as well as the early adult stage of *C. sulstoni.* From these data, we confirmed the expression of *C. sulstoni lin-4, mir-48, mir-84, mir-241*, and *mir-x*, and found that their developmental dynamics in *C. sulstoni* are similar to *C. elegans* and *C. macrosperma* (Fig. 3A–3C and S4A-S4C; Table S1).

To assess the potential functional conservation of protein coding components of the heterochronic pathway, we used RNAi to knock down heterochronic gene homologs in *C. sulstoni.* Previous studies reported that loss-of-function mutations of *lin-14, lin-28*, or *hbl-1* in *C. elegans* result in precocious alae formation (Ambros and Horvitz, 1984; Ambros, 1989; Fay et al., 1999; Abrahante et al., 2003; Lin et al., 2003). Similarly, in *C. sulstoni* RNAi of *lin-14, lin-28*, or *hbl-1* caused essentially identical phenotypes (Fig. 4A and 4B) as previously reported for *C. elegans*.

Previous studies also reported that *lin-46(lf)* results in a mild retarded phenotype manifesting as minor gaps and branches in adult alae (Pepper et al., 2004), and *lin-29(lf)* results in a more severe retarded phenotype manifesting as significant gaps in or complete absence of adult alae (Ambros and Horvitz, 1984; Ambros, 1989). RNAi of *lin-46* or *lin-29* in *C. sulstoni* resulted in nearly identical phenotypes (Fig. 4C–4E) as previously reported for *C. elegans lin-46(lf)* or *lin-29(lf)* mutants.

The similarities in expression of *lin-4* and the *let-7-family*, the conservation of complementary sites in the 3’ UTRs of *lin-14, lin-28*, and *hbl-1*, and the RNAi knock down phenotypes for *lin-14, lin-28, lin-46, hbl-1*, and *lin-29* indicate that the heterochronic pathway is largely conserved between *C. sulstoni* and *C. elegans.*

### Temporal regulation and function of LIN-41 are largely conserved in *C. sulstoni*

MicroRNA families are groups of microRNAs that share an identical seed sequence (nucleotides 2-8) but differ in their non-seed sequence (Roush and Slack, 2008). In principle, members of the same microRNA family can regulate the same target via seed pairing, but at the same time, differences in non-seed sequences can allow family member specificity of targeting through specific microRNA base-pairing with non-seed nucleotides (Moore et al., 2015; Broughton et al., 2016; Brancati and Grosshans, 2018). In *C. elegans*, the *let-7-family* consists of *let-7, mir-48, mir-84, mir-241*, and *mir-795* (Table S1) (Abbott et al., 2005; Roush and Slack, 2008). Interestingly, *C. elegans let-7* targets the *lin-41* 3’ UTR via two sites that are “weakly” complementary to the *let-7-family* seed sequence (hereafter referred to as SM for “seed match”), both of which are supplemented with significant non-seed pairings to *let-7*. This base pairing configuration (hereafter referred to as SM+SUP) is thought to confer specificity for regulation of *lin-41* by *let-7*, to the exclusion of the other *let-7-family* microRNAs. Interestingly, the “weak” seed of each SM is distinct from the other: pairing of *let-7* to the first SM results in a bulged target adenine in the seed helix between the g4 and g5 (hereafter referred to as the “bulge-SM”; Fig. 5A); for the second SM, the seed helix contains a G-U wobble base pair at g5 (hereafter referred to as the “GU-SM”; Fig. 5B) (Reinhart et al., 2000; Vella et al., 2004; Ecsedi et al., 2015).

**Figure 5.**
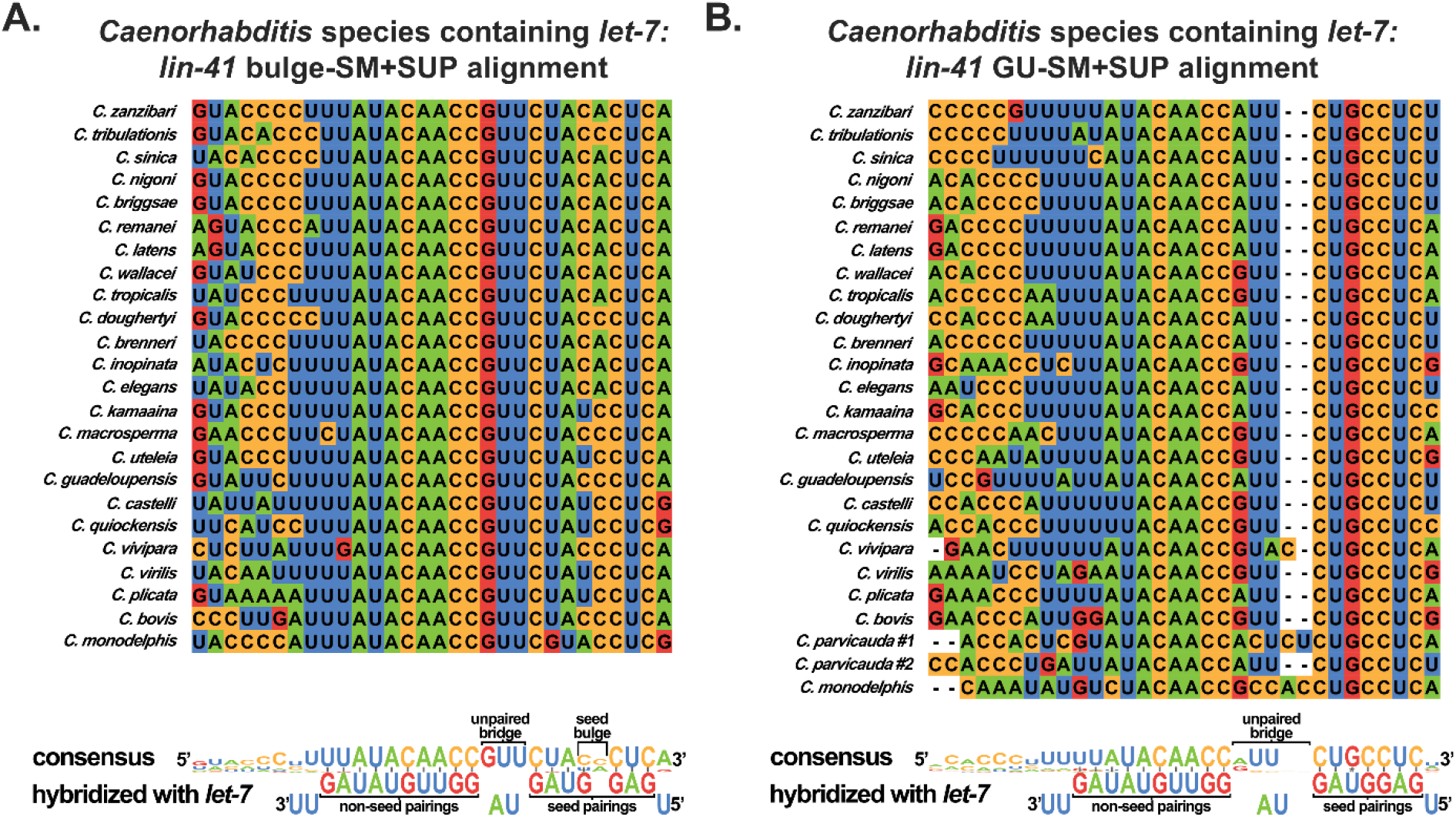
The *let-7* seed match (SM) plus supplemental non-seed pairing (+SUP) sites in the 3’ UTR of *lin-41* are highly conserved in *Caenorhabditis* species that contain *let-7*. Sequence alignment of the bulge-SM+SUP site (A) and the GU-SM+SUP (B) in the 3’ UTR of *lin-41* in *Caenorhabditis* species containing *let-7.* Shown at the bottom of each panel is the respective consensus sequence predicted hybridization with *let-7*. Watson-Crick base pairing is shown with a solid line between the paired bases. The G-U base pair in the seed is shown with an asterisk between the paired bases.

To gauge the phylogenetic conservation of the *let-7* SM+SUP configurations in the *lin-41* 3’ UTR regions of other related species, we aligned predicted *lin-41* SM+SUP sequences for all available *Caenorhabditis* species genomes. To identify potential *let-7* family base paring SM+SUP sequences, we searched downstream of the stop codon of *lin-41* homologs for sequences complementary to the *let-7* seed sequence, including bulges and G-U pairings. With the exception of the species lacking *let-7*, all *Caenorhabditis* species have two “weak” SM+SUPs in their *lin-41* 3’ UTR regions, and apart from *C. parvicauda*, which has two GU-SM+SUPs, all *let-7*-containing *Caenorhabditis* species have one bulge-SM+SUP positioned 5’ of one GU-SM+SUP (Fig. 5A and 5B; Tables S3 and S4). Moreover, the SM+SUPs are always in relatively close proximity to each other, with the 3’ most base-paired nucleotide of the bulge-SM being no further than 39 nucleotides away from the 5’ most base-paired nucleotide of the GU-SM (Tables S3 and S4). These results indicate that there is strong selective pressure to maintain a strict *let-7*-specific regulation of *lin-41* in *Caenorhabditis* species that express *let-7* microRNA.

One possible explanation for how the absence of *let-7* is accommodated in *Japonica* species is that one or more of the other *let-7-family* microRNAs may have adopted the role of regulating *lin-41.* To test this possibility, we examined the *lin-41* 3’ UTR regions of *let-7*-lacking genomes for sequences complementary to the remaining *let-7-family* microRNAs. Remarkably, in all seven of these genomes, we observed conservation of the GU-SM, indicating conservation of the regulation of *lin-41* by one or more *let-7-family* microRNAs, despite the absence of *let-7* itself (Fig. 6A and 6B; Table S4). Interestingly, we failed to observe any significant conservation of *lin-41* 3’ UTR sequences adjacent to the GU-SM, arguing against any conservation of supplemental non-seed pairing by a particular *let-7-family* microRNA (Fig. 6A and 6B; Table S4).

**Figure 6.**
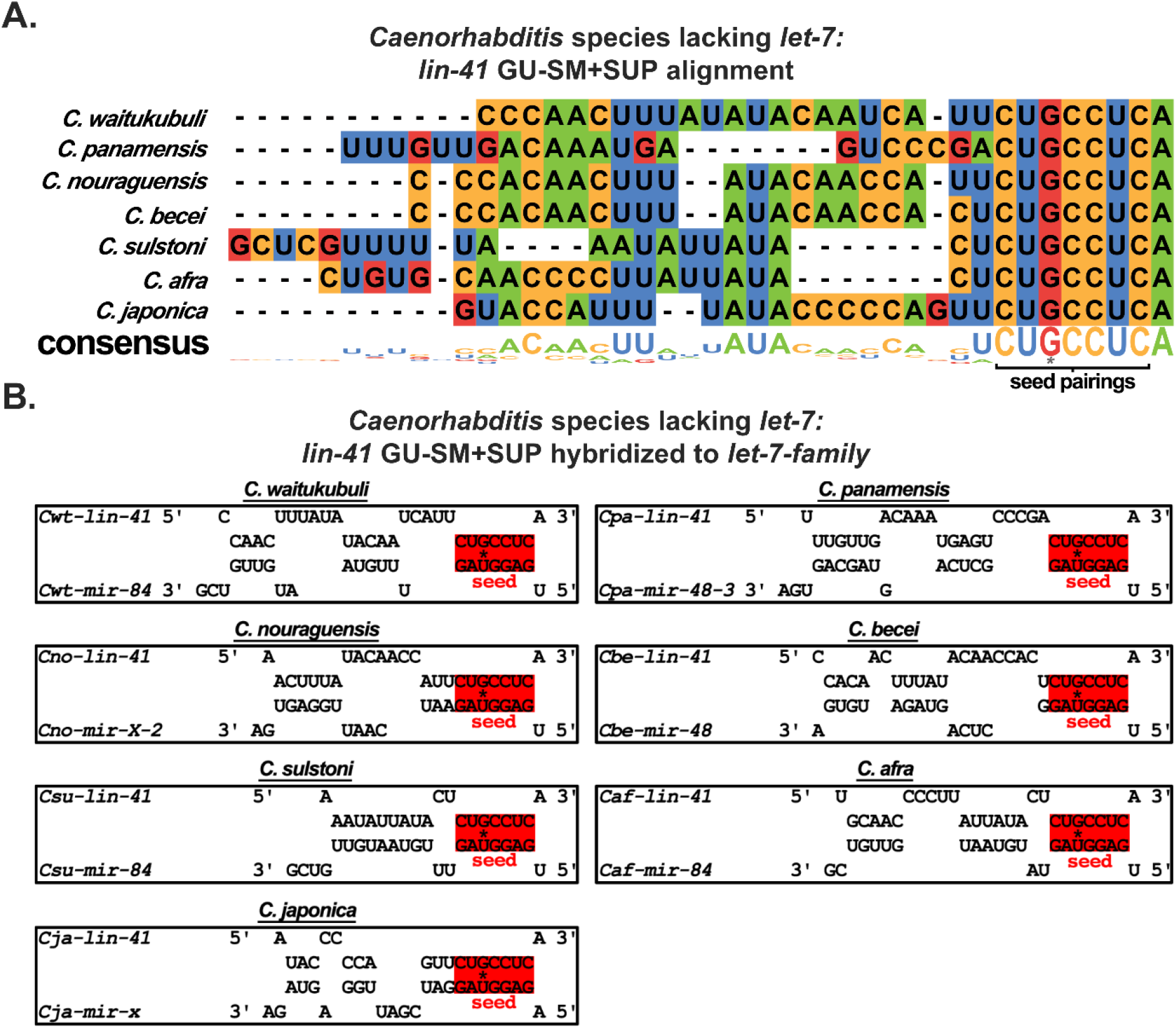
The seed match (SM) plus supplemental non-seed pairing (+SUP) site in the 3’ UTR of *lin-41* are divergent amongst *Caenorhabditis* species lacking *let-7*. (A) Sequence alignment of the GU-SM+SUP in the 3’ UTR of *lin-41* in *Caenorhabditis* species lacking *let-7.* The G-U base pair in the seed pairings is shown with an asterisk. (B) Predicted RNA hybridizations of *let-7-family* microRNAs with the GU-SM+SUP in the *lin-41* 3’ UTR regions in *Caenorhabditis* species lacking *let-7.* Shown are the hybridizations of the *let-7-family* microRNA with the most favorable hybridization (lowest MFE) for each respective species. The G-U base pair in the seed pairings is shown with an asterisk.

Owing to the lack of conservation of supplemental non-seed matching sequence in the *lin-41* 3’UTR of *Caenorhabditis* species lacking *let-7*, we hypothesized that one or more *let-7-family* microRNAs could regulate *lin-41* via GU-SM+SUP base-pairing pattern, but the particular *let-7*-family microRNAs with the best match to *lin-41* could vary for each species. We presumed that for a *let-7-family* microRNA to assume a similar role as *let-7* in regulating *lin-41*, the minimum free energy (MFE) between that *let-7-family* microRNA and the *lin-41* GU-SM+SUP should be similar to that for the interaction of *let-7* with *lin-41* in *let-7*-containing species. To test this, we calculated the MFE of base pairing for each *let-7*-lacking species’ predicted *let-7-family* microRNAs to their respective *lin-41* GU-SM+SUP and compared it to the MFEs of the *let-7-family* microRNAs hybridized to the bulge-SM+SUP as well as GU-SM+SUP in all *let-7*-containing species. In all *let-7*-containing species, the lowest (most favorable) MFE pairing of a *let-7-family* microRNA to the *lin-41* bulge-SM+SUP and GU-SM+SUP was for the interaction with *let-7* (average MFE of −27.2 ± 0.5 kcal/mol and −29.2 ± 1.2 kcal/mol, respectfully; Fig. S7A and S7B; Tables S3 and S4). By contrast, in *let-7*-lacking species, the *let-7-family* microRNA that had the lowest MFE pairing to the *lin-41* GU-SM+SUP varied between species and was higher (less favorable) than the MFE of *let-7* with either the bulge-SM+SUP or GU-SM+SUP in species that contain *let-7* (average MFE −21.5 ± 2.1 kcal/mol; Fig. S7B; Table S4).

LIN-41 is an RNA binding protein that, in *C. elegans*, forms distinct foci in the cytoplasm particularly around the periphery of the nucleus (Fig. 7A) (Spike et al., 2014). In the hypodermis of *C. elegans*, LIN-41 is expressed during the L1 through L3 stages and is undetectable in the L4 and adult stages due to its translational repression by *let-7* (Fig. 7A) (Slack et al., 2000). The relatively weaker predicted MFE of the interaction between *let-7-family* microRNAs and *lin-41* in species lacking *let-7* compared to *let-7* and *lin-41* in species containing *let-7* suggests that the *let-7-family* could have a less prominent role in down regulation of *lin-41* in species lacking *let-7* than *let-7* does in species containing *let-7*. To determine whether LIN-41 protein is down regulated in a representative *let-7*-lacking species as observed in *C. elegans*, we used CRISPR to GFP-tag the N-terminus of endogenous LIN-41 in *C. sulstoni*. We observed a nearly identical expression pattern in *C. sulstoni* to what we observed in *C. elegans*: hypodermal LIN-41 is expressed in the L1 through L3 stages and is downregulated in the L4 and adult stages (Fig. 7A and 7B). This down regulation of LIN-41 is indicative of prominent temporal regulation similar to what is observed in *C. elegans* and could reflect the action of one or more *let-7-family* microRNAs.

**Figure 7.**
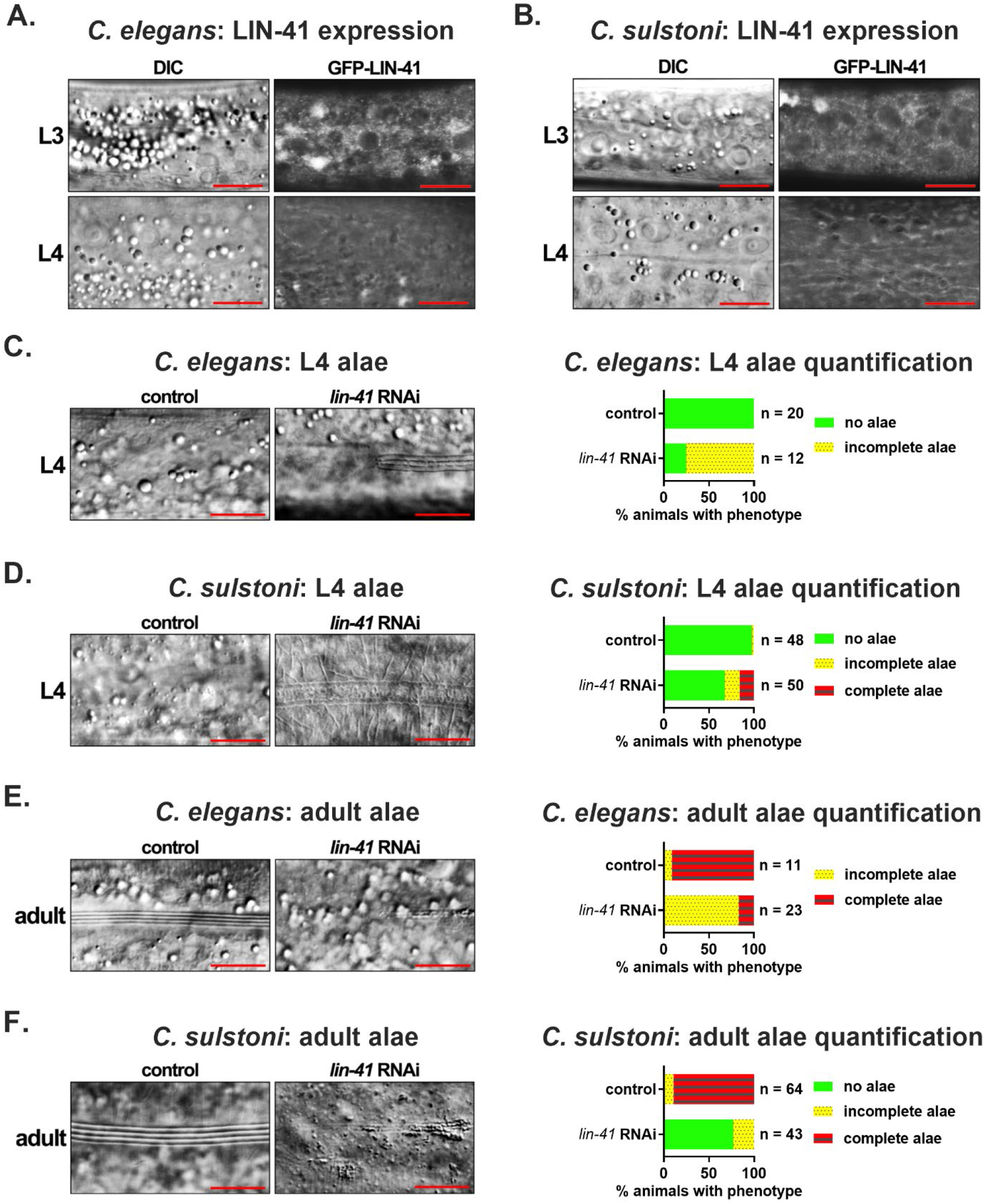
*lin-41* expression and function are conserved between *C. elegans* and *C. sulstoni*. (A and B) Representative DIC images (left panels) and endogenously tagged GFP-LIN-41 images (right panels) of hypodermal cells in *C. elegans* (A) and *C. sulstoni* (B) L3 (top panels) and L4 animals (bottom panels). Scale bars = 10 μM. (C-F) Representative DIC images of *C. elegans* L4 hypodermis (C), *C. sulstoni* L4 hypodermis (D), *C. elegans* adult hypodermis (E), and *C. sulstoni* adult hypodermis (F) of animals fed control (empty vector) (left panels) and *lin-41* RNAi (middle panels) and quantification of alae phenotypes (graphs). Scale bars = 10 μM. Note: RNAi experiments used for Figures 4B, 4D, 7D and 7F were performed together. Therefore, control RNAi data used for Figure 4B and 4D were also used for Figure 7D and 7F.

In *C. elegans, lin-41(lf)* animals precociously produce adult alae during the L4 stage (Slack et al., 2000). To determine if LIN-41 could be functionally conserved in species that lack *let-7*, we used RNAi to knock down *lin-41* in *C. sulstoni.* Similar to *C. elegans*, when *lin-41* was knocked down in *C. sulstoni* we observed L4 animals with adult alae. However, precocious alae occurred in a lower percentage of animals than previously reported for *lin-41(lf)* in *C. elegans*, and generally the precocious adult alae were of rather indistinct morphology compared to bona fide adult alae (Fig. 7D and 7E) (Reinhart et al., 2000; Slack et al., 2000; Banerjee et al., 2005; Nolde et al., 2007). The relatively low penetrance of the precocious alae phenotype could possibly reflect poor RNAi knock down of *lin-41*. To test this, we knocked down *lin-41* in our GFP-tagged strain and confirmed efficient knockdown by an absence of detectable GFP-LIN-41 fluorescence in hypodermal cells. Moreover, animals in which *lin-41* was knocked down were sterile, a hallmark of *lin-41* loss-of-function for its germline function in *C. elegans* (data not shown) (Slack et al., 2000).

To our surprise, RNAi of *lin-41* in *C. sulstoni* caused not only precocious and morphologically abnormal alae at the L4 stage, but also caused a highly penetrant alae formation phenotype in adult stage animals, where alae were often absent, incomplete, and/or morphologically abnormal (Fig. 7F). This sort of adult stage phenotype had not been reported previously for *lin-41(lf)* in *C. elegans.* The previous studies of *lin-41(lf)* in *C. elegans* employed partial loss-of-function mutations and not the high efficiency RNAi expression vector that we used to knock down *lin-41* in *C. sulstoni* (T444T vector) (Sturm et al., 2018). We hypothesized that the functions of *lin-41* could be conserved between *C. elegans* and *C. sulstoni*, and the discrepancies in the phenotypes previously reported and observed here could be due to the partial function of the *lin-41* alleles used in previous studies. To test this hypothesis, we knocked down *lin-41* in *C. elegans* using the new, high-efficiency vector (T444T), and we observed similar phenotypes observed in *C. sulstoni*: L4 larvae with adult alae and incomplete/weak alae in adults (Fig. 7C–7F). Interestingly, the L4 alae phenotypes observed in *C. sulstoni* were still less penetrant and weaker in appearance than what we observed in *C. elegans* (Fig. 7C and 7D). Moreover, the alae phenotypes observed in *C. elegans* adults appeared weaker than what we observed in *C. sulstoni* (Fig. 7E and 7F). Aside from these differences, our results indicate that the temporal patterning and function of *lin-41* are largely conserved between *C. elegans* and *C. sulstoni*.

## DISCUSSION

*let-7* appears to be indispensable across diverse bilaterian phyla, indicating deep and pervasive evolutionary constraints on maintaining the entire 22-nt sequence of *let-7*. Moreover, *let-7*’s function in promoting cellular differentiation and repression of pluripotency are conserved as well, which could reflect some degree of conservation of orthologous targets. Chiefly among these are *lin-41* (*TRIM71* in mammals) and *lin-28* (*LIN28* in mammals) both of which are pluripotency-promoting RNA binding proteins whose expression is directly repressed by *let-7* in invertebrates and vertebrates alike (Slack et al., 2000; Kloosterman et al., 2004; Schulman et al., 2005; Lin et al., 2007; Vella et al., 2004; Ding and Grosshans, 2009; Ecsedi et al., 2015). Consequently, loss-of-function of *let-7* results in dysregulation of developmental progression (Reinhart et al., 2000; Kloosterman et al., 2004; Sokol et al., 2008; Caygill and Johnston, 2008), and can lead to disease such as cancer (Balzeau et al., 2017). Thus, *let-7* is an important regulator of animal development and tissue homeostasis.

In this study, we characterized for the first time a cohort of animal species lacking the *let-7* microRNA sequence in their genomes. Using microRNA profiling, we confirmed the absence of mature *let-7* expression, while also confirming the conserved expression of the *let-7-family* microRNAs *mir-48, mir-84*, and *mir-241.* We also found that, except for the absence of *let-7*, the heterochronic pathway, into which *let-7* is deeply integrated in *C. elegans*, appears otherwise functionally conserved in a representative *let-7*-lacking species, *C. sulstoni*. Finally, we provide evidence that *lin-41*, a *let-7*-specific target in *C. elegans*, is regulated by other members of the *let-7-family* of microRNAs in *Caenorhabditis* species that lack *let-7*.

Our finding that *C. macrosperma* appears to be an exceptional member of the *Japonica* group that contains *let-7* raises interesting questions. Based on the current phylogenetic tree of *Caenorhabditis* species, for *C. macrosperma* to retain *let-7*, while the remaining members of the *Japonica* group lack *let-7*, three possibilities are suggested: 1) *C. macrosperma* does not belong in the *Japonica* group, 2) *C. macrosperma* is correctly assigned to the *Japonica* group, and gained the *let-7* sequence after it was lost in a *Japonica* ancestor, or 3) *C. macrosperma* is correctly assigned to *Japonica*, and *let-7* was lost at least three times during the evolution of the *Japonica* group.

The evidence strongly suggests that *C. macrosperma* is indeed correctly placed in the *Japonica* clade. Previous publications have routinely recovered *C. macrosperma* as a member of *Japonica* group with the most recent publications being a highly extensive, genome-wide studies with high statistical likelihood of *C. macrosperma*’s correct recovery (Kiontke et al., 2011; Felix et al., 2014; Stevens et al., 2019; Stevens et al., 2020). Our results further strengthen this argument. We found that in *C. macrosperma*, the protein sequence of the *let-7*-specific target *lin-41* was most similar to *lin-41* protein sequence in other *Japonica* group species, and homologs of the novel *let-7-family* microRNA *mir-x* that we identified in *C. macrosperma* can only be found in *Japonica* group species. Put together, these suggests that *C. macrosperma*’s recovery in the *Japonica* group is correct.

We can imagine two possible ways by which *C. macrosperma* could have gained *let-7*, 1) in a spontaneous manner, possibly by random sequence drift of a *let-7-family* paralog, or 2) through gene transfer from another *Caenorhabditis.* The first scenario is unlikely because the *let-7* genomic region in *C. macrosperma* is syntenic to the *let-7* region in *C. elegans*, arguing against spontaneous gain as no other *Japonica* group member has a *let-7-family* paralogue in that syntenic region. The second scenario also seems unlikely, as there are no known examples of horizontal gene transfer between *Caenorhabditis* species. Moreover, the presence in *C. macrosperma* of *lin-41* bulge-SM+SUP and GU-SM+SUP sites matching *let-7* that are nearly identical to the *lin-41* bulge-SM+SUP and GU-SM+SUP sites in every other *let-7*-containing *Caenorhabditis* species strongly suggests that *C. macrosperma* inherited *let-7*, and its targeting of *lin-41*, from a common ancestor of the other *Japonica* species, and that *let-7* was lost during the evolution of other *Japonica* species.

A parsimonious model for loss of *let-7* in the *Japonica* clade suggests at least three independent loss events – once in the common ancestor to *C. waitukubuli, C. panamensis, C. nouraguensis*, and *C. becei*, a second loss in a common ancestor to *C. sulstoni* and *C. afra*, and a third loss in *C. japonica.* Multiple independent losses of *let-7* in the *Japonica* group, together with the dramatically lower expression level of *let-7* in *C. macrosperma* compared to the relatively robust expression of *let-7* in *C. elegans*, suggests that *let-7* became relatively dispensable in a common ancestor of these *Japonica* species. We suggest that a hallmark of the evolution of the *Japonica* group may be a reduced dependency on *let-7* for functions that are otherwise critical in most other *Caenorhabditis* species.

The extensive synteny of the *let-7* region of *C. elegans* with *C. sulstoni* suggest that loss of *let-7* was not associated with dramatic genome rearrangements in the region. Only one gene neighboring *let-7* in *C. elegans, C05G5.7*, also appears to have been fully lost in *C. sulstoni.* The loss of this gene might suggest a functional relationship between *let-7* and *C05G5.7*, perhaps even that loss of *C05G5.7* could functionally compensate for loss of *let-7.* Interestingly, *C05G5.7* is transcribed on the same primary transcript as *let-7* and serves as a negative regulator of *let-7* processing. However, *C05G5.7* appears not to be conserved outside of *C. elegans*, and loss off *C05G5.7* fails to suppress *let-7* loss-of-function (Nelson and Ambros, 2019).

A key component of the heterochronic network in *C. elegans* is the RNA binding protein LIN-41. In *C. elegans*, loss of *let-7* is lethal, primarily due to the de-repression of *lin-41* (Ecsedi et al., 2015; Aeschimann et al., 2019). During wild type *C. elegans* development, LIN-41 levels decrease during the mid-to-late-L4 stage due to a sharp increase in *let-7* expression and subsequent *let-7*-mediated inhibition of LIN-41 translation. *let-7* interacts with the 3’ UTR of *lin-41* mRNA via base pairing to two non-canonical seed match (SM) sites – one with a bulge adenosine of the *lin-41* mRNA in the seed helix between the g4 and g5 and the other with a G-U wobble base pair at the g5 – combined with extensive supplemental (SUP) non-seed pairings. In *Caenorhabditis* species that contain *let-7*, both SM+SUPs are highly conserved. In the set of *Caenorhabditis* species that lack *let-7*, the *lin-41* 3’ UTR contains only the GU-SM, with no extensive conservation of non-seed pairings.

To be sure, the presence of a *let-7-family* seed match in the 3’ UTR of *lin-41* in species lacking *let-7* indicates that *lin-41* is likely regulated by one or more of the remaining *let-7*-*family* microRNAs. However, several considerations suggest that *let-7-family* mediated regulation of *lin-41* in these species may be relatively less critical compared to *let-7*-containing species, where the extensive conservation of two SM+SUP sites indicates that both of these sites are necessary for robust regulation of *lin-41* by *let-7.* By contrast, the *lin-41* 3’ UTRs of *Caenorhabditis* species lacking *let-7* contain just one *let-7-family* SM site of relatively weak predicted binding (higher MFE) and do not exhibit evidence of conserved non-seed pairing.

In species lacking *let-7*, the *let-7*-*family* microRNA(s) predicted to regulate *lin-41* via SM+SUP pairing varied between species. *mir-84* was most favorable in three species (*C. waitukubuli, C. sulstoni*, and *C. afra*), *mir-48* was most favorable in two species (*C. panamensis* and *C. becei*), and *mir-x* was the most favorable in two species (*C. nouraguensis* and *C. japonica*) (Fig. 6B; Table S4). Moreover, in some species that lack *let-7*, the *let-7-family* microRNA with the most favorable MFE was not much better than the next best MFE. For example, in *C. panamensis, mir-48* was the most favorable with an MFE of −25.8 kcal/mol, while *mir-84* was the next best with an MFE of −25.2 kcal/mol. This variability in which *let-7-family* microRNA is most favorable in combination with the relatively close MFEs of multiple *let-7-family* microRNAs within a species suggests that the regulation of *lin-41* by the *let-7-family* is less constrained, in terms of ortholog specificity, in *let-7*-lacking species than in *let-7*-containing species.

We found that despite the lack of *let-7* in *C. sulstoni* the temporal expression pattern and function of LIN-41 protein is largely conserved; LIN-41 levels decrease during the L4 stage of *C. sulstoni*, as is the case in *C. elegans*, and knockdown of *lin-41* results in similar heterochronic phenotypes in *C. sulstoni* and *C. elegans.* While our findings indicate that one or more *let-7-family* microRNA(s) may stand in for the absent *let-7* to developmentally regulate *C. sulstoni lin-41*, we cannot rule out contributions from transcriptional or other post-transcriptional mechanisms.

In this study, we have identified a cohort of related *Caenorhabditis* species within the *Japonica* group that lack the highly conserved *let-7* microRNA. Our findings that one member of the *Japonica* group, *C. macrosperma*, has retained *let-7*, together with the pattern of species affinities within the *Japonica* group, suggests that *let-7* was lost at least three times during the evolution of the *Japonica* clade. We do not currently know what allowed for the loss of *let-7* in these species. However, it appears that in the *Japonica* group, *let-7* was seemingly released from its evolutionary-entrenched regulatory relationship with *lin-41*, thereby allowing for the regulation of *lin-41* to be taken over by the *let-7-family*, and this hypothetical evolutionary release of *let-7* apparently resulted in the dispensability of *let-7.* Further study is required to determine the nature of evolutionary factors that could cause an otherwise deeply conserved microRNA such as *let-7* to become dispensable.

## MATERIALS AND METHODS

### Nematode methods

All *Caenorhabditis* species were cultured on nematode growth medium (NGM) (Brenner, 1974) and fed with *E. coli* HB101 except for all RNAi experiments, in which *C. elegans* and *C. sulstoni* were fed *E. coli* HT115. Synchronized populations of developmentally staged worms were obtained using standard methods (Stiernagle, 2006). All experiments involving *C. elegans*, unless otherwise noted, were performed at 20°C. All experiments with the other *Caenorhabditis* species, unless otherwise noted, were performed at 25°C. A list of strains used in this study is in Table S5.

### Staging, developmental, and phenotypic analyses of *C. elegans, C. sulstoni*, and *C. macrosperma*

To characterize the effects of temperature on development, synchronized populations of *C. elegans* and *C. sulstoni* were plated at 15°C, 20°C, 25°C, 30°C, 33°C, and 35°C. A synchronized population of *C. macrosperma* was plated at 25°C. Except for *C. sulstoni* animals plated at 15°C, developing populations plated at each respective temperature were observed every hour until animal development reached the adult stage. Following the initial 48 hours of hourly observation, *C. sulstoni* animals plated at 15°C were observed every 12 hours until animals reached the adult stage. L1 alae, lethargy, cuticular molting, gonad migration, vulva development, adult alae, and oogenesis were used as markers to determine and calibrate developmental stages.

For heterochronic phenotype analyses, larvae were fed with RNAi food (as described below) starting from the L1 stage, and subsequent animals of defined developmental stages (as described above) were picked from healthy uncrowded mixed staged cultures and imaged. DIC microscopy was used to image hypodermis and alae. Fluorescence microscopy was used to image GFP-LIN-41.

For quantification of alae formation, the entire length of the animal’s cuticle was observed using DIC microscopy. Alae with one or more discontinuity was scored as incomplete. Any region where alae branched into multiple directions was scored as a branch.

### Microscopy

All DIC and fluorescent images were obtained using a ZEISS Imager Z1 equipped with ZEISS Axiocam 503 mono camera, and the ZEN Blue software. Prior to imaging, worms were anesthetized with 0.2 mM levamisole in M9 buffer and mounted on 2% agarose pads. Adobe Photoshop was used to adjust the brightness and contrast of the images to enhance the visualization of the DIC and fluorescent signals. All fluorescent images were taken using the same microscopy settings and a standard exposure time for each larval stage for each reporter (*C. elegans* GFP-LIN-41 and *C. sulstoni* GFP-LIN-41). Identical brightness and contrast adjustments were used for each fluorescent image.

### *Caenorhabditis* genomes

All genomes used in this study were provided by the *Caenorhabditis* Genomes Project (http://caenorhabditis.org).

### Syntenic comparisons

Syntenic comparisons were performed using GEvo (https://genomevolution.org/coge/GEvo.pl) (Lyons and Freeling, 2008) with the following algorithm settings – Alignment Algorithm: (B)LastZ: Large Regions; Word size: 8; Gap start penalty: 300; Gap extend penalty: 30; Chaining: chain and extend; Score threshold: 2000; Mask threshold: 0; Minimum HSP length for finding overlapped features: 50.

### Identification of homologous genes

Identification of all homologs were performed using CoGeBlast and the *Caenorhabditis* Genomes Project Blast webpages (https://genomevolution.org/coge/CoGeBlast.pl; http://blast.caenorhabditis.org/).

### Sequence Alignments

Sequence alignments were performed using Clustal Omega (www.ebi.ac.uk) (Madeira et al., 2019) and visualized using Jalview (www.jalview.org) (Waterhouse et al., 2009).

### RNA hybridization predictions and MFE calculations

RNA hybridization predictions and MFE calculations were performed using RNAhybrid (https://bibiserv2.cebitec.uni-bielefeld.de) (Rehmsmeier et al., 2004).

### *lin-41* phylogeny

*lin-41* phylogenetic tree was generated from orthology clustering provided by the *Caenorhabditis* Genomes Project (Stevens, 2020) and visualized using iTOL (https://itol.embl.de/) (Letunic and Bork, 2019). The *lin-41* phylogenetic tree was rooted on the outgroup *Diploscapter coronatus*.

### RNA extraction

Populations of animals were collected and flash-frozen in liquid nitrogen, and total RNA was extracted using Qiazol reagen (Qiagen) as described by (McJunkin and Ambros, 2017).

### FirePlex microRNA detection

microRNAs were quantified from total RNA using FirePlex miRNA assay (Abcam) following the manufacturer’s instructions. Guava easyCyte 8HT (Millipore) was used for analysis.

### Small RNA sequencing and *let-7-family* microRNA identification and normalization

Samples of total RNA were used to generate all small RNA libraries using a QIAseq miRNA Library Kit (Qiagen) following the manufacturer’s instructions. All libraries were sequenced on an Illumina NextSeq 500 sequencer.

*let-7-family* microRNAs were identified by searching for reads that contained the *let-7-family* seed sequence “GAGGTAG” at positions 2-8. *let-7-family* microRNA reads were considered legitimate if the sequence mapped to the genome and if the read was predicted to form a microRNA-like stem-loop secondary structure with adjacent genomic sequence. RNA secondary structure modeling was performed using RNAfold (http://rna.tbi.univie.ac.at/cgi-bin/RNAWebSuite/RNAfold.cgi) (Lorenz et al., 2011). In most instances, reads corresponding to microRNA precursors were identified, adding additional credence to the validation of *let-7-family* microRNAs.

microRNAs were quantified by normalizing the number of a given microRNA reads per million total reads in that library.

### GFP tagging of *C. sulstoni* LIN-41

*C. sulstoni GFP-lin-41* was generated using CRISPR/Cas9 methods adapted from Paix et al. and Dokshin et al. (Paix et al., 2014; Paix et al., 2015; Dokshin et al., 2018). The germlines of young adult females were injected with a mix of CRISPR RNA (crRNA) that targeted the 5’ end of the *lin-41* coding sequence and the ‘co-CRISPR’ marker *dpy-10*, tracrRNA (Table S6), a PCR-derived dsDNA HR template, Cas9 protein (IDT) and IDT’s duplex buffer (30 mM HEPES, pH 7.5; 100 mM potassium acetate). L4 females from plates where F1 animals that exhibited the co-CRISPR phenotype were picked, mated to a single male picked from the same plate, allowed to lay eggs, and then genotyped using PCR. F2s with GFP expression were cloned from F1s that scored positively by PCR genotyping for the desired modification. Single male and single female progeny were then mated, and a homozygous line was selected by GFP expression and PCR genotyping and subjected to Sanger sequencing for validation. The mutant was then thrice backcrossed to wild type. Sequence details of the *GFP::lin-41* allele generated in this study can be found in Table S7.

### Bacterial RNAi feeding strain constructions

cDNA from mixed staged *C. elegans* and *C. sulstoni* was generated from total RNA using SuperScript IV Reverse Transcriptase (ThermoFisher) and oligoDT following the manufacturer’s instructions. PCR was then used to amplify portions of *C. elegans lin-41, C. sulstoni lin-14, C. sulstoni lin-28, C. sulstoni lin-29, C. sulstoni lin-41, C. sulstoni lin-46, C. sulstoni hbl-1*, and *C. sulstoni unc-22*, respectively. Primers (Table S6) used for each respective PCR also added KpnI sites to each end of each PCR product except for *C. elegans lin-41*, which added a HindIII site to one end and a KpnI site to the other end. The PCR products and the T444T vector were digested with KpnI (NEB) restriction enzyme for all *C. sulstoni* genes and HindIII and KpnI for *C. elegans lin-41*. The cut T444T vector was then dephosphorylated, and the cut PCR products and the cut/dephosphorylated vector were gel purified, ligated, and transformed into TOP10 chemically competent cells. Purified plasmids were subjected to Sanger sequencing for validation and transformed into chemically competent *E. coli* HT115 cells.

### RNAi knockdown of heterochronic genes

RNAi by feeding *C. elegans* and *C. sulstoni* with the strains described above was conducted as described in Conte et al. (Conte et al., 2015).

## Data availability

All *Caenorhabditis* strains will be available at the *Caenorhabditis* Genetics Center (https://cgc.umn.edu). Reagents used in this study are available upon request. Raw small RNA sequencing data will be deposited in the NCBI SRA (http://www.ncbi.nlm.nih.gov/sra). Normalized small RNA sequencing data used for Figures 3 and S4 can be found in Table S8. The exact timing data used for Figures 3, S3, and S4 can be found in Table S9. The RNAi quantification data used for Figures 4 and 7 can be found in Table S10.

## ACKNOWLEDGEMENTS

We thank the members of the Ambros and the Mello laboratories for helpful discussions and comments on this project, especially Ye Duan for help with small RNA sequencing. We thank Lewis Stevens and Mark Blaxter for help with the *Caneorhabditis* phylogeny and *lin-41* gene tree. We thank the *Caenorhabditis* Genomes project for providing the *Caenorhabditis* genomic sequences and annotations. Some strains were provided by the *Caenorhabditis* Genetics Center (CGC), which is funded by NIH Office of Research Infrastructure Programs (P40 OD010440). Resources for the genomic analyses were provided by CoGe, which is funded by the NSF (IOS-1444490). The T444T plasmid was a gift from Tibor Vellai (Addgene plasmid # 113081; http://n2t.net/addgene:113081; RRID:Addgene_113081).

## AUTHOR CONTRIBUTIONS

Conceptualization: C.N., V.A.; Methodology: C.N., V.A.; Formal analysis: C.N., V.A.; Investigation: C.N.; Resources: V.A.; Data curation: C.N.; Writing - original draft: C.N.; Writing - review & editing: C.N., V.A.; Supervision: V.A.; Project administration: V.A.; Funding acquisition: V.A.

## REFERENCES

Abbott, A. L., Alvarez-Saavedra, E., Miska, E. A., Lau, N. C., Bartel, D. P., Horvitz, H. R. and Ambros, V. (2005). The let-7 MicroRNA family members mir-48, mir-84, and mir-241 function together to regulate developmental timing in Caenorhabditis elegans. Dev Cell 9, 403–14.

Abrahante, J. E., Daul, A. L., Li, M., Volk, M. L., Tennessen, J. M., Miller, E. A. and Rougvie, A. E. (2003). The Caenorhabditis elegans hunchback-like gene lin-57/hbl-1 controls developmental time and is regulated by microRNAs. Dev Cell 4, 625–37.

Aeschimann, F., Neagu, A., Rausch, M. and Grosshans, H. (2019). let-7 coordinates the transition to adulthood through a single primary and four secondary targets. Life Sci Alliance 2.

Ambros, V. (1989). A hierarchy of regulatory genes controls a larva-to-adult developmental switch in C. elegans. Cell 57, 49–57.

Ambros, V. and Horvitz, H. R. (1984). Heterochronic mutants of the nematode Caenorhabditis elegans. Science 226, 409–16.

Ambros, V. and Ruvkun, G. (2018). Recent Molecular Genetic Explorations of Caenorhabditis elegans MicroRNAs. Genetics 209, 651–673.

Balzeau, J., Menezes, M. R., Cao, S. and Hagan, J. P. (2017). The LIN28/let-7 Pathway in Cancer. Front Genet 8, 31.

Banerjee, D., Kwok, A., Lin, S. Y. and Slack, F. J. (2005). Developmental timing in C. elegans is regulated by kin-20 and tim-1, homologs of core circadian clock genes. Dev Cell 8, 287–95.

Bartel, D. P. (2009). MicroRNAs: target recognition and regulatory functions. Cell 136, 215–33.

Brancati, G. and Grosshans, H. (2018). An interplay of miRNA abundance and target site architecture determines miRNA activity and specificity. Nucleic Acids Res 46, 3259–3269.

Brenner, S. (1974). The genetics of Caenorhabditis elegans. Genetics 77, 71–94.

Broughton, J. P., Lovci, M. T., Huang, J. L., Yeo, G. W. and Pasquinelli, A. E. (2016). Pairing beyond the Seed Supports MicroRNA Targeting Specificity. Mol Cell 64, 320–333.

Caygill, E. E. and Johnston, L. A. (2008). Temporal regulation of metamorphic processes in Drosophila by the let-7 and miR-125 heterochronic microRNAs. Curr Biol 18, 943–50.

Chalfie, M., Horvitz, H. R. and Sulston, J. E. (1981). Mutations that lead to reiterations in the cell lineages of C. elegans. Cell 24, 59–69.

Conte, D., Jr., Macneil, L. T., Walhout, A. J. M. and Mello, C. C. (2015). RNA Interference in Caenorhabditis elegans. Curr Protoc Mol Biol 109, 26 3 1–26 3 30.

De Wit, E., Linsen, S. E., Cuppen, E. and Berezikov, E. (2009). Repertoire and evolution of miRNA genes in four divergent nematode species. Genome Res 19, 2064–74.

Ding, X. C. and Grosshans, H. (2009). Repression of C. elegans microRNA targets at the initiation level of translation requires GW182 proteins. EMBO J28, 213–22.

Dokshin, G. A., Ghanta, K. S., Piscopo, K. M. and Mello, C. C. (2018). Robust Genome Editing with Short Single-Stranded and Long, Partially Single-Stranded DNA Donors in Caenorhabditis elegans. Genetics 210, 781–787.

Ecsedi, M., Rausch, M. and Grosshans, H. (2015). The let-7 microRNA directs vulval development through a single target. Dev Cell 32, 335–44.

Fay, D. S., Stanley, H. M., Han, M. and Wood, W. B. (1999). A Caenorhabditis elegans homologue of hunchback is required for late stages of development but not early embryonic patterning. Dev Biol 205, 240–53.

Felix, M. A., Braendle, C. and Cutter, A. D. (2014). A streamlined system for species diagnosis in Caenorhabditis (Nematoda: Rhabditidae) with name designations for 15 distinct biological species. PLoS One 9, e94723.

Heo, I., Joo, C., Cho, J., Ha, M., Han, J. and Kim, V. N. (2008). Lin28 mediates the terminal uridylation of let-7 precursor MicroRNA. Mol Cell 32, 276–84.

Ilbay, O. and Ambros, V. (2019). Regulation of nuclear-cytoplasmic partitioning by the lin-28-lin-46 pathway reinforces microRNA repression of HBL-1 to confer robust cell-fate progression in C. elegans. Development 146.

Kiontke, K. C., Felix, M. A., Ailion, M., Rockman, M. V., Braendle, C., Penigault, J. B. and Fitch, D. H. (2011). A phylogeny and molecular barcodes for Caenorhabditis, with numerous new species from rotting fruits. BMC Evol Biol 11, 339.

Kloosterman, W. P., Wienholds, E., Ketting, R. F. and Plasterk, R. H. (2004). Substrate requirements for let-7 function in the developing zebrafish embryo. Nucleic Acids Res 32, 6284–91.

Lau, N. C., Lim, L. P., Weinstein, E. G. and Bartel, D. P. (2001). An abundant class of tiny RNAs with probable regulatory roles in Caenorhabditis elegans. Science 294, 858–62.

Lee, R. C., Feinbaum, R. L. and Ambros, V. (1993). The C. elegans heterochronic gene lin-4 encodes small RNAs with antisense complementarity to lin-14. Cell 75, 843–54.

Lehrbach, N. J., Armisen, J., Lightfoot, H. L., Murfitt, K. J., Bugaut, A., Balasubramanian, S. and Miska, E. A. (2009). LIN-28 and the poly(U) polymerase PUP-2 regulate let-7 microRNA processing in Caenorhabditis elegans. Nat Struct Mol Biol 16, 1016–20.

Letunic, I. and Bork, P. (2019). Interactive Tree Of Life (iTOL) v4: recent updates and new developments. Nucleic Acids Res 47, W256–W259.

Lim, L. P., Lau, N. C., Weinstein, E. G., Abdelhakim, A., Yekta, S., Rhoades, M. W., Burge, C. B. and Bartel, D. P. (2003). The microRNAs of Caenorhabditis elegans. Genes Dev 17, 991–1008.

Lin, S. Y., Johnson, S. M., Abraham, M., Vella, M. C., Pasquinelli, A., Gamberi, C., Gottlieb, E. and Slack, F. J. (2003). The C elegans hunchback homolog, hbl-1, controls temporal patterning and is a probable microRNA target. Dev Cell 4, 639–50.

Lin, Y. C., Hsieh, L. C., Kuo, M. W., Yu, J., Kuo, H. H., Lo, W. L., Lin, R. J., Yu, A. L. and Li, W. H. (2007). Human TRIM71 and its nematode homologue are targets of let-7 microRNA and its zebrafish orthologue is essential for development. Mol Biol Evol 24, 2525–34.

Lorenz, R., Bernhart, S. H., Honer Zu Siederdissen, C., Tafer, H., Flamm, C., Stadler, P. F. and Hofacker, I. L. (2011). ViennaRNA Package 2.0. Algorithms Mol Biol 6, 26.

Lyons, E. and Freeling, M. (2008). How to usefully compare homologous plant genes and chromosomes as DNA sequences. Plant J 53, 661–73.

Madeira, F., Park, Y. M., Lee, J., Buso, N., Gur, T., Madhusoodanan, N., Basutkar, P., Tivey, A. R. N., Potter, S. C., Finn, R. D., et al. (2019). The EMBL-EBI search and sequence analysis tools APIs in 2019. Nucleic Acids Res 47, W636–W641.

Mcjunkin, K. and Ambros, V. (2017). A microRNA family exerts maternal control on sex determination in C. elegans. Genes Dev 31, 422–437.

Moore, M. J., Scheel, T. K., Luna, J. M., Park, C. Y., Fak, J. J., Nishiuchi, E., Rice, C. M. and Darnell, R. B. (2015). miRNA-target chimeras reveal miRNA 3’-end pairing as a major determinant of Argonaute target specificity. Nat Commun 6, 8864.

Moss, E. G., Lee, R. C. and Ambros, V. (1997). The cold shock domain protein LIN-28 controls developmental timing in C. elegans and is regulated by the lin-4 RNA. Cell 88, 637–46.

Nam, Y., Chen, C., Gregory, R. I., Chou, J. J. and Sliz, P. (2011). Molecular basis for interaction of let-7 microRNAs with Lin28. Cell 147, 1080–91.

Nelson, C. and Ambros, V. (2019). Trans-splicing of the C. elegans let-7 primary transcript developmentally regulates let-7 microRNA biogenesis and let-7 family microRNA activity. Development 146.

Newman, M. A., Thomson, J. M. and Hammond, S. M. (2008). Lin-28 interaction with the Let-7 precursor loop mediates regulated microRNA processing. RNA 14, 1539–49.

Nolde, M. J., Saka, N., Reinert, K. L. and Slack, F. J. (2007). The Caenorhabditis elegans pumilio homolog, puf-9, is required for the 3’UTR-mediated repression of the let-7 microRNA target gene, hbl-1. Dev Biol 305, 551–63.

Paix, A., Folkmann, A., Rasoloson, D. and Seydoux, G. (2015). High Efficiency, Homology-Directed Genome Editing in Caenorhabditis elegans Using CRISPR-Cas9 Ribonucleoprotein Complexes. Genetics 201, 47–54.

Paix, A., Wang, Y., Smith, H. E., Lee, C. Y., Calidas, D., Lu, T., Smith, J., Schmidt, H., Krause, M. W. and Seydoux, G. (2014). Scalable and versatile genome editing using linear DNAs with microhomology to Cas9 Sites in Caenorhabditis elegans. Genetics 198, 1347–56.

Pasquinelli, A. E., Reinhart, B. J., Slack, F., Martindale, M. Q., Kuroda, M. I., Maller, B., Hayward, D. C., Ball, E. E., Degnan, B., Muller, P., et al. (2000). Conservation of the sequence and temporal expression of let-7 heterochronic regulatory RNA. Nature 408, 86–9.

Pepper, A. S., Mccane, J. E., Kemper, K., Yeung, D. A., Lee, R. C., Ambros, V. and Moss, E. G. (2004). The C. elegans heterochronic gene lin-46 affects developmental timing at two larval stages and encodes a relative of the scaffolding protein gephyrin. Development 131, 2049–59.

Piskounova, E., Polytarchou, C., Thornton, J. E., Lapierre, R. J., Pothoulakis, C., Hagan, J. P., Iliopoulos, D. and Gregory, R. I. (2011). Lin28A and Lin28B inhibit let-7 microRNA biogenesis by distinct mechanisms. Cell 147, 1066–79.

Rehmsmeier, M., Steffen, P., Hochsmann, M. and Giegerich, R. (2004). Fast and effective prediction of microRNA/target duplexes. RNA 10, 1507–17.

Reinhart, B. J., Slack, F. J., Basson, M., Pasquinelli, A. E., Bettinger, J. C., Rougvie, A. E., Horvitz, H. R. and Ruvkun, G. (2000). The 21-nucleotide let-7 RNA regulates developmental timing in Caenorhabditis elegans. Nature 403, 901–6.

Roush, S. and Slack, F. J. (2008). The let-7 family of microRNAs. Trends Cell Biol 18, 505–16.

Ruby, J. G., Jan, C., Player, C., Axtell, M. J., Lee, W., Nusbaum, C., Ge, H. and Bartel, D. P. (2006). Large-scale sequencing reveals 21U-RNAs and additional microRNAs and endogenous siRNAs in C. elegans. Cell 127, 1193–207.

Rybak, A., Fuchs, H., Smirnova, L., Brandt, C., Pohl, E. E., Nitsch, R. and Wulczyn, F. G. (2008). A feedback loop comprising lin-28 and let-7 controls pre-let-7 maturation during neural stem-cell commitment. Nat Cell Biol 10, 987–93.

Schulman, B. R., Esquela-Kerscher, A. and Slack, F. J. (2005). Reciprocal expression of lin-41 and the microRNAs let-7 and mir-125 during mouse embryogenesis. Dev Dyn 234, 1046–54.

Shi, Z., Montgomery, T. A., Qi, Y. and Ruvkun, G. (2013). High-throughput sequencing reveals extraordinary fluidity of miRNA, piRNA, and siRNA pathways in nematodes. Genome Res 23, 497–508.

Slack, F. J., Basson, M., Liu, Z., Ambros, V., Horvitz, H. R. and Ruvkun, G. (2000). The lin-41 RBCC gene acts in the C. elegans heterochronic pathway between the let-7 regulatory RNA and the LIN-29 transcription factor. Mol Cell 5, 659–69.

Sokol, N. S., Xu, P., Jan, Y. N. and Ambros, V. (2008). Drosophila let-7 microRNA is required for remodeling of the neuromusculature during metamorphosis. Genes Dev 22, 1591–6.

Spike, C. A., Coetzee, D., Eichten, C., Wang, X., Hansen, D. and Greenstein, D. (2014). The TRIM-NHL protein LIN-41 and the OMA RNA-binding proteins antagonistically control the prophase-to-metaphase transition and growth of Caenorhabditis elegans oocytes. Genetics 198, 1535–58.

Stefani, G., Chen, X., Zhao, H. and Slack, F. J. (2015). A novel mechanism of LIN-28 regulation of let-7 microRNA expression revealed by in vivo HITS-CLIP in C. elegans. RNA 21, 985–96.

Stevens, L. (2020). CGP orthology clustering. Zenodo. https://doi.org/10.5281/zenodo.4068211

Stevens, L., Felix, M. A., Beltran, T., Braendle, C., Caurcel, C., Fausett, S., Fitch, D., Frezal, L., Gosse, C., Kaur, T., et al. (2019). Comparative genomics of 10 new Caenorhabditis species. Evol Lett 3, 217–236.

Stevens, L., Rooke, S., Falzon, L. C., Machuka, E. M., Momanyi, K., Murungi, M. K., Njoroge, S. M., Odinga, C. O., Ogendo, A., Ogola, J., et al. (2020). The Genome of Caenorhabditis bovis. Curr Biol 30, 1023–1031 e4.

Stiernagle, T. (2006). Maintenance of C. elegans. WormBook, 1–11.

Stratoulias, V., Heino, T. I. and Michon, F. (2014). Lin-28 regulates oogenesis and muscle formation in Drosophila melanogaster. PLoS One 9, e101141.

Sturm, A., Saskoi, E., Tibor, K., Weinhardt, N. and Vellai, T. (2018). Highly efficient RNAi and Cas9-based auto-cloning systems for C. elegans research. Nucleic Acids Res 46, e105.

Sulston, J. E. and Horvitz, H. R. (1977). Post-embryonic cell lineages of the nematode, Caenorhabditis elegans. Dev Biol 56, 110–56.

Tsialikas, J., Romens, M. A., Abbott, A. and Moss, E. G. (2017). Stage-Specific Timing of the microRNA Regulation of lin-28 by the Heterochronic Gene lin-14 in Caenorhabditis elegans. Genetics 205, 251–262.

Van Wynsberghe, P. M., Kai, Z. S., Massirer, K. B., Burton, V. H., Yeo, G. W. and Pasquinelli, A. E. (2011). LIN-28 co-transcriptionally binds primary let-7 to regulate miRNA maturation in Caenorhabditis elegans. Nat Struct Mol Biol 18, 302–8.

Vella, M. C., Choi, E. Y., Lin, S. Y., Reinert, K. and Slack, F. J. (2004). The C. elegans microRNA let-7 binds to imperfect let-7 complementary sites from the lin-41 3’UTR. Genes Dev 18, 132–7.

Viswanathan, S. R., Daley, G. Q. and Gregory, R. I. (2008). Selective blockade of microRNA processing by Lin28. Science 320, 97–100.

Waterhouse, A. M., Procter, J. B., Martin, D. M., Clamp, M. and Barton, G. J. (2009). Jalview Version 2--a multiple sequence alignment editor and analysis workbench. Bioinformatics 25, 1189–91.

