## supplemental tables and figures for "A cohort of *Caenorhabditis* species lacking the highly conserved *let-7* microRNA"

| Species | *let-7* sequence | *mir-48* sequence(s) | *mir-84* sequence(s) | *mir-241* sequence | *mir-795* sequence | *mir-x* sequence(s) | *lin-4* sequence(s) |
|---|---|---|---|---|---|---|---|
| *C. tribulationis* | TGAGGTAGTAGGTTGTATAGTT | TGAGGTAGGCTCAGTAGATGCGA | TGAGGTAGTTTGTAATGCTGTCG | TGAGGTAGGTGCGAGAAATGA | no *mir-795* | no *mir-x* | TCCCTGAGACCTCAAGTGTGA |
| *C. zanzibari* | TGAGGTAGTAGGTTGTATAGTT | TGAGGTAGGCTCAGTAGATGCGA | TGAGGTAGTTTGTAATGTTGTG | TGAGGTAGGTGCGAGAAATGA | no *mir-795* | no *mir-x* | TCCCTGAGACCTCAAGTGTGA |
| *C. sinica* | TGAGGTAGTAGGTTGTATAGTT | TGAGGTAGGCTCAGTAGATGCGA | TGAGGTAGTTTGTAATGCTGTG | TGAGGTAGGTGCGAGAAATGA | no *mir-795* | no *mir-x* | TCCCTGAGACCTCAAGTGTGA |
| *C. nigoni* | TGAGGTAGTAGGTTGTATAGTT | TGAGGTAGGCTCAGTAGATGCGA | TGAGGTAGTTTGCAATGCTGTCG | TGAGGTAGGTGCGAGAAATGA | no *mir-795* | no *mir-x* | TCCCTGAGACCTCAAGTGTGA |
| *C. briggsae* | TGAGGTAGTAGGTTGTATAGTT[2] | TGAGGTAGGCTCAGTAGATGCGA[2] | TGAGGTAGTTTGCAATGCTGTC[2] | TGAGGTAGGTGTGAGAAATGA[2] | no *mir-795* | no *mir-x* | TCCCTGAGACCTCAAGTGTGA[2] |
| *C. remanei* | TGAGGTAGTAGGTTGTATAGTT[2] | TGAGGTAGGCTCAGTAGATGCG[2] | TGAGGTAGTTTGTAATGCTGTCG[2] | AGAGGTAGGTGCGAGAAATG[2] | no *mir-795* | no *mir-x* | TCCCTGAGACCTCAAGTGTGA[2] |
| *C. latens* | TGAGGTAGTAGGTTGTATAGTT | TGAGGTAGGCTCAGTAGATGCGA<br>(2 copies) | TGAGGTAGTTTGTAATGCTGTCG | AGAGGTAGGTGCGAGAAATGA<br>(2 copies) | no *mir-795* | no *mir-x* | TCCCTGAGACCTCAAGTGTGA |
| *C. brenneri* | TGAGGTAGTAGGTTGTATAGTT[2] | TGAGGTAGGCTCAGTAGATGCGA[2]<br>(2 copies) | TGAGGTAGTTTGGAAATGCTGTCG[2] | AGAGGTAGGTGCGAGAAATGA[2]<br>(2 copies) | no *mir-795* | no *mir-x* | TCCCTGAGACCTCAAGTGTGA[2]<br>(2 copies) |
| *C. wallacei* | TGAGGTAGTAGGTTGTATAGTT | TGAGGTAGGCTCAGTAGATGCGA | TGAGGTAGTTTGTAATGCTGTCG | TGAGGTAGGTGCGAGAAATGA | no *mir-795* | no *mir-x* | TCCCTGAGACCTCAAGTGTGA |
| *C. tropicalis* | TGAGGTAGTAGGTTGTATAGTT | TGAGGTAGGCTCAGTAGATGCGA | TGAGGTAGTTTGTAATGCTGTCG | TGAGGTAGGTGCGAGAAATGA | no *mir-795* | no *mir-x* | TCCCTGAGACCTCAAGTGTGA |
| *C. doughertyi* | TGAGGTAGTAGGTTGTATAGTT | TGAGGTAGGCTCAGTAGATGCGA<br>(2 copies) | TGAGGTAGTTTGTAATGCTGTCG | TGAGGTAGGTGCGGGAAAGG<br>TGAGGTAGATGTGAGAAATGA<br>(2 copies) | no *mir-795* | no *mir-x* | TCCCTGAGACCTCAAGTGTGA |
| *C. inopinata* | TGAGGTAGTAGGTTGTATAGTT[1] | TGAGGTAGGCTCAGTAGATGCGA[1] | TGAGGTAGTTTGAAATGCTGTCG | TGAGGTAGGTGCGAGAAATGA[1] | no *mir-795* | no *mir-x* | TCCCTGAGACCTCAAGTGTGA[1] |
| *C. elegans* | TGAGGTAGTAGGTTGTATAGTT[1,2,3] | TGAGGTAGGCTCAGTAGATGCGA[1,2,3] | TGAGGTAGTATGTAATATTGTAGA[1,2,3] | TGAGGTAGGTGCGAGAAATGA[1,2,3] | TGAGGTAGATTGATCAGCGAGCTT[1,2,3] | no *mir-x* | TCCCTGAGACCTCAAGTGTGA[1,2,3] |
| *C. kamaaina* | TGAGGTAGTAGGTTGTATAGTT[1] | TGAGGTAGGCTCAGTAGATGTGA[1] | TGAGGTAGTTTGTAATGTTGTCG | TGAGGTAGGTGCGAGAAATGA[1] | no *mir-795* | no *mir-x* | TCCCTGAGACCTCAAGTGTGA[1] |
| *C. waitukubuli* | no *let-7* | TGAGGTAGGCTCAGTAGATGTGA[1] | TGAGGTAGTTTGTAATGTTGTCG[#] | TGAGGTAGGTGCGAGAAATGA[1] | no *mir-795* | TGAGGTAGAATCAATTGCAGTGA[#] | TCCCTGAGACCTCAAGTGTGA[1]<br>(2 copies) |
| *C. panamensis* | no *let-7* | TGAGGTAGGCTCAGTAGATGTGA[1*]<br>TGAGGTAGGCTCAGTAGTTGTGA[1*]<br>TGAGGTAGGCTCAGTAGCAGTGA[1*]<br>TGAGGTAGGCTCAGTGACGTGA[1*] | TGAGGTAGTTTGCAATGTTGTCG[#] | TGAGGTAGGTGCGAGAAATGA[1*]<br>TGAGGTAGCGTGCGAGAAATGA[1,*,#] | no *mir-795* | TGAGGTAGGATCAATTGCAGTGA[#] | TCCCTGAGACCTCAAGTGTGA[1] |
| *C. nouraguensis* | no *let-7* | TGAGGTAGGCTCAGTAGATGTGA[1] | TGAGGTAGTTTGCAATGTTGTCG[*,#]<br>(2 copies)<br>TGAGGTAGTTTGCAATGTTGTAG[*,#] | TGAGGTAGGTGCGAGAAATGA[1] | no *mir-795* | TGAGGTAGGATCAATTGGAGTGA[#]<br>TGAGGTAGAATCAATTGGAGTGA[#]<br>TGAGGTAGAAGCAATTGCAGTGA[#] | TCCCTGAGACCTCAAGTGTGA[1] |
| *C. becei* | no *let-7* | TGAGGTAGGCTCAGTAGATGTGA[1] | TGAGGTAGTTTGCAATGTTGTCG[#]<br>(3 copies) | TGAGGTAGGTGCGAGAAATGA[1] | no *mir-795* | TGAGGTAGAATCAATTGCAGTGA[#]<br>TGAGGTAGAATCAATTGGAGTGA[#] | TCCCTGAGACCTCAAGTGTGA[1] |
| *C. macrosperma* | TGAGGTAGTAGGTTGTATAGTT[3] | TGAGGTAGGCTCAGTAGATGTGA[1,3] | TGAGGTAGTTTGCAATGTTGTCG[3,#] | TGAGGTAGGTGCGAGAAATGA[1,3] | no *mir-795* | TGAGGTAGAATCAATTGCAGTGA[3,#] | TCCCTGAGACCTCAAGTGTGA[1,3] |
| *C. sulstoni* | no *let-7* | TGAGGTAGGCTCAGTAGATGTGA[1,3] | TGAGGTAGTTTGTAATGTTGTCG[3,#] | TGAGGTAGGTGCGAGAAATG[1,3] | no *mir-795* | TGAGGTAGGATCAAATTGAGTGA[3,#] | TCCCTGAGACCTCAAGTGTGA[1,3] |
| *C. afra* | no *let-7* | TGAGGTAGGCTCAGTAGATGTGA[1] | TGAGGTAGTATGTAATGTTGTCG[#] | TGAGGTAGGTGCGAGAAATG[1] | no *mir-795* | TGAGGTAGGATCCAACCGAGCGA[#] | TCCCTGAGACCTCAAGTGTGA[1] |
| *C. japonica* | no *let-7* | TGAGGTAGGCTTTGTAGATGTGA[1] | TGAGGTAGTATTTAATGTTGTTG[#] | TGAGGTAGGTGCGAGAAATGA[1] | no *mir-795* | AGAGGTAGGATCGATTGGAGTAGA[#] | TCCCTGAGACCTCAAGTGTGA[1] |
| *C. guadeloupensis* | TGAGGTAGTAGGTTGTATAGTT | no *mir-48* | TGAGGTAGTTTGTAAAGTTGTCG | no *mir-241* | no *mir-795* | no *mir-x* | TCCCTGAGACCTCAAGTGTGA |
| *C. uteleia* | TGAGGTAGTAGGTTGTATAGTT | no *mir-48* | TGAGGTAGTTTGTAATGTTGTAG | no *mir-241* | no *mir-795* | no *mir-x* | TCCCTGAGACCTCAAGTGTGA |
| *C. quiockensis* | TGAGGTAGTAGGTTGTATAGTT | no *mir-48* | TGAGGTAGTTTAAAATGTTGTCG | no *mir-241* | no *mir-795* | no *mir-x* | TCCCTGAGACCTCAAGTGTGA<br>TCCCTGAGTCCTCAAATGTGA |
| *C. castelli* | TGAGGTAGTAGGTTGTATAGTT | no *mir-48* | TGAGGTAGTTTAAAATGTTGTCG | no *mir-241* | no *mir-795* | no *mir-x* | TCCCTGAGACCTCAAGTGTGA |
| *C. angaria* | TGAGGTAGTAGGTTGTATAGTT | no *mir-48* | TGAGGTAGTTTAAAATGTTGTCG | no *mir-241* | no *mir-795* | no *mir-x* | TCCCTGAGACCTCAAGTGTGA<br>(2 copies) |
| *C. vivipara* | TGAGGTAGTAGGTTGTATAGTT | no *mir-48* | TGAGGTAGTTTAAAATGTTGTCG | no *mir-241* | no *mir-795* | no *mir-x* | TCCCTGAGACCTCAAGTGTGA |
| *C. virilis* | TGAGGTAGTAGGTTGTATAGTT | no *mir-48* | TGAGGTAGTTTGAAATGTTGTCG | no *mir-241* | no *mir-795* | no *mir-x* | TCCCTGAGACCTCAAGTGTGA |
| *C. plicata* | TGAGGTAGTAGGTTGTATAGTT | no *mir-48* | TGAGGTAGTTTTCGATGTTGTCG | no *mir-241* | no *mir-795* | no *mir-x* | TCCCTGAGACCTCAAGTGTGA |
| *C. bovis* | AGAGGTAGTAGGTTGTATAGTT | no *mir-48* | TGAGGTAGTTTTAAATGTTGTCG | no *mir-241* | no *mir-795* | no *mir-x* | TCCCTGAGACCTCAAGTGTGA |
| *C. parvicauda* | AGAGGTAGTAGGTTGTATAGTT | no *mir-48* | no *mir-84* | no *mir-241* | no *mir-795* | no *mir-x* | TCCCTGAGACCTCAAGTGTGA |
| *C. monodelphis* | TGAGGTAGTAGGTTGTATAGTT | no *mir-48* | TGAGGTAGTATTAAATGTTGTCG | no *mir-241* | no *mir-795* | no *mir-x* | TCCCTGAGACCTCAAGTGCGA |

**Table S1. *let-7-family* and *lin-4* microRNA sequences in *Caenorhabditis* species.**

List of all predicted and confirmed *let-7-family* and *lin-4* microRNA sequences in all *Caenorhabditis* species used in this study. Outlined in the green box are species recovered in the *Elegans* group. Outlined in the red box are species recovered in the *Japonica* group. [1]confirmed using FirePlex assay (this study). [2]confirmed in previous study (Pasquinelli et al., 2000; Reinhart et al., 2000; Lau et al., 2001; Lim et al., 2003; Ruby et al., 2006; de Wit et al., 2009; Shi et al., 2013). [3]confirmed using small RNA sequencing (this study). [*]unable to distinguish which copy was detected in FirePlex assay. [#]likely in *mir-84* polycistron.

| Species | Gene | Gene ID | 3' UTR length (nt) | Position (nt) of *lin-4* complementary site(s) after stop codon | Position (nt) of *let-7-family* complementary site(s) after stop codon |
|---|---|---|---|---|---|
| ***C. elegans*** | *lin-14* | WBGene00003003 | 1604 | 838, 886, 1117 | 429, 882, 896, 1295 |
| ***C. sulstoni*** | *lin-14* | CSP32.g2871 | unknown | 974, 1021, 1263 | 556, 1017, 1031 |
| ***C. elegans*** | *lin-28* | WBGene00003014 | 531 | 341 | 321 |
| ***C. sulstoni*** | *lin-28* | CSP32.g6943 | unknown | 145 | 128 |
| ***C. elegans*** | *lin-29* | WBGene00003015 | 697 | none | none |
| ***C. sulstoni*** | *lin-29* | CSP32.g16252 | unknown | none | none |
| ***C. elegans*** | *lin-41* | WBGene00003026 | 1165 | none | 709, 757 |
| ***C. sulstoni*** | *lin-41* | CSP32.g4866 | unknown | none | 655 |
| ***C. elegans*** | *lin-46* | WBGene00003031 | 153 | none | none |
| ***C. sulstoni*** | *lin-46* | CSP32.g17530 | unknown | none | none |
| ***C. elegans*** | *hbl-1* | WBGene00001824 | 1394 | none | 272, 940, 1210, 1253 |
| ***C. sulstoni*** | *hbl-1* | CSP32.g179 | unknown | none | 240, 621, 818, 934, 1156, 1198 |

**Table S2. *C. sulstoni* homologs of *C. elegans* heterochronic genes.**

List of the *C. sulstoni* CDS name, its *C. elegans* homolog of all heterochronic genes tested in RNAi experiments for this study, the *C. elegans* 3’ UTR lengths for each respective heterochronic gene, and the position after the stop codon of *lin-4* and *let-7-family* seed complementary site(s) in both *C. elegans* and *C. sulstoni.*

| **Species** | **Bulge-SM+SUP sequence** | **Distance (nt) from stop codon** | **MFE (kcal/mol) with *let-7*** | **MFE (kcal/mol) with *mir-48*** | **MFE (kcal/mol) with *mir-84*** | **MFE (kcal/mol) with *mir-241*** | **MFE (kcal/mol) with *mir-795*** | **MFE (kcal/mol) with *mir-x*** |
|---|---|---|---|---|---|---|---|---|
| *C. zanzibari* | GUACCCCUUUAUACAACCGUUCUACACUCA | 520 | **-27.2** | -14.6 | -17.7 | -15.9 | no *mir-795* | no *mir-x* |
| *C. tribulationis* | GUACACCCUUAUACAACCGUUCUACCCUCA | 657 | **-27.1** | -14.6 | -18.8 | -16.2 | no *mir-795* | no *mir-x* |
| *C. sinica* | UACACCCCUUAUACAACCGUUCUACACUCA | 720 | **-27.1** | -14.6 | -16.1 | -16.2 | no *mir-795* | no *mir-x* |
| *C. nigoni* | GUACCCCUUUAUACAACCGUUCUACACUCA | 488 | **-27.2** | -14.6 | -14.6 | -15.9 | no *mir-795* | no *mir-x* |
| *C. briggsae* | GUACCCCUUUAUACAACCGUUCUACACUCA | 458 | **-27.2** | -14.6 | -14.6 | -15.9 | no *mir-795* | no *mir-x* |
| *C. remanei* | AGUACCCAUUAUACAACCGUUCUACACUCA | 771 | **-28.2** | -14.9 | -16.2 | -16.0 | no *mir-795* | no *mir-x* |
| *C. latens* | AGUACCCUUUAUACAACCGUUCUACACUCA | 758 | **-27.2** | -14.7 | -16.1 | -16.0 | no *mir-795* | no *mir-x* |
| *C. wallacei* | GUAUCCCUUUAUACAACCGUUCUACACUCA | 863 | **-27.2** | -17.2 | -16.2 | -16.2 | no *mir-795* | no *mir-x* |
| *C. tropicalis* | UAUCCCUUUUAUACAACCGUUCUACACUCA | 864 | **-27.2** | -14.6 | -16.1 | -16.2 | no *mir-795* | no *mir-x* |
| *C. doughertyi* | GUACCCCCUUAUACAACCGUUCUACCCUCA | 612 | **-27.1** | -13.9 | -16.1 | -17.7 & -21.6 | no *mir-795* | no *mir-x* |
| *C. brenneri* | UACCCCUUUUAUACAACCGUUCUACACUCA | 884 | **-27.2** | -14.6 | -17.9 | -16.0 | no *mir-795* | no *mir-x* |
| *C. inopinata* | AUACUCUUUUAUACAACCGUUCUACACUCA | 807 | **-27.2** | -14.6 | -14.4 | -16.2 | no *mir-795* | no *mir-x* |
| *C. elegans* | UAUACCUUUUAUACAACCGUUCUACACUCA | 709 | **-27.2** | -14.6 | -17.1 | -16.2 | -19.0 | no *mir-x* |
| *C. kamaaina* | GUACCCUUUUAUACAACCGUUCUAUCCUCA | 724 | **-27.2** | -15.8 | -17.8 | -16.2 | no *mir-795* | no *mir-x* |
| *C. waitukubuli* | No Bulge-SM | not available | No *let-7* | not available | not available | not available | no *mir-795* | not available |
| *C. panamensis* | No Bulge-SM | not available | No *let-7* | not available | not available | not available | no *mir-795* | not available |
| *C. nouraguensis* | No Bulge-SM | not available | No *let-7* | not available | not available | not available | no *mir-795* | not available |
| *C. becei* | No Bulge-SM | not available | No *let-7* | not available | not available | not available | no *mir-795* | not available |
| *C. macrosperma* | GAACCCUUCUAUACAACCGUUCUACCCUCA | 697 | **-28.6** | -16.0 | -18.0 | -16.2 | no *mir-795* | -16.9 |
| *C. sulstoni* | No Bulge-SM | not available | No *let-7* | not available | not available | not available | no *mir-795* | not available |
| *C. afra* | No Bulge-SM | not available | No *let-7* | not available | not available | not available | no *mir-795* | not available |
| *C. japonica* | No Bulge-SM | not available | No *let-7* | not available | not available | not available | no *mir-795* | not available |
| *C. uteleia* | GUACCCUUUUAUACAACCGUUCUAUCCUCA | 792 | **-27.2** | no *mir-48* | -17.7 | no *mir-241* | no *mir-795* | no *mir-x* |
| *C. guadeloupensis* | GUAUUCUUUUAUACAACCGUUCUACCCUCA | 608 | **-27.2** | no *mir-48* | -17.1 | no *mir-241* | no *mir-795* | no *mir-x* |
| *C. castelli* | UAUUAUUUUUAUACAACCGUUCUAUCCUCG | 426 | **-26.8** | no *mir-48* | -17.2 | no *mir-241* | no *mir-795* | no *mir-x* |
| *C. angaria* | Unable to identify due to sequence gaps | not available | not available | no *mir-48* | not available | no *mir-241* | no *mir-795* | no *mir-x* |
| *C. quiockensis* | UUCAUCCUUUAUACAACCGUUCUAUCCUCG | 399 | **-26.8** | no *mir-48* | -17.2 | no *mir-241* | no *mir-795* | no *mir-x* |
| *C. vivipara* | CUCUUAUUUGAUACAACCGUUCUACCCUCA | 933 | **-25.6** | no *mir-48* | -17.6 | no *mir-241* | no *mir-795* | no *mir-x* |
| *C. virilis* | UACAAUUUUUAUACAACCGUUCUAUCCUCA | 637 | **-27.2** | no *mir-48* | -12.4 | no *mir-241* | no *mir-795* | no *mir-x* |
| *C. plicata* | GUAAAAAUUUAUACAACCGUUCUAUCCUCA | 706 | **-27.2** | no *mir-48* | -17.8 | no *mir-241* | no *mir-795* | no *mir-x* |
| *C. bovis* | CCCUUGAUUUAUACAACCGUUCUACCCUCA | 968 | **-27.0** | no *mir-48* | -17.6 | no *mir-241* | no *mir-795* | no *mir-x* |
| *C. parvicauda* | No Bulge-SM | not available | not available | no *mir-48* | no *mir-84* | no *mir-241* | no *mir-795* | no *mir-x* |
| *C. monodelphis* | UACCCCAUUUAUACAACCGUUCGUACCUCG | 2257 | **-26.8** | no *mir-48* | -18.0 | no *mir-241* | no *mir-795* | no *mir-x* |

**Table S3. *lin-41* bulge-SM+SUP sequences and MFEs with all *let-7-family* microRNAs.**
List of all *lin-41* bulge-SM+SUP in all *Caenorhabditis* species used in this study. The position of the bulge-SM relative to the stop codon, and all predicted MFEs (kcal/mol) of each bulge-SM+SUP sequence hybridized with each respective *let-7-family* microRNA. MFEs in bold = lowest energy (most favorable). Outlined in the green box are species recovered in the *Elegans* group. Outlined in the red box are species recovered in the *Japonica* group.

| Species | GU-SM+SUP sequence(s) | Distance (nt) from stop codon | MFE (kcal/mol) with *let-7* | MFE (kcal/mol) with *mir-48* | MFE (kcal/mol) with *mir-84* | MFE (kcal/mol) with *mir-241* | MFE (kcal/mol) with *mir-795* | MFE (kcal/mol) with *mir-x* |
|---|---|---|---|---|---|---|---|---|
| *C. zanzibari* | CCCCCGUUUUUAUACAACCAUUCUGCCUCU | 575 | **-29.0** | -20.0 | -20.6 | -20.4 | no *mir-795* | no *mir-x* |
| *C. tribulationis* | CCCCCUUUUAUAUACAACCAUUCUGCCUCU | 710 | **-28.7** | -16.9 | -21.7 | -16.9 | no *mir-795* | no *mir-x* |
| *C. sinica* | CCCCUUUUUUCAUACAACCAUUCUGCCUCU | 774 | **-27.3** | -17.3 | -19.0 | -19.0 | no *mir-795* | no *mir-x* |
| *C. nigoni* | ACACCCCUUUUAUACAACCAUUCUGCCUCU | 521 | **-29.0** | -16.9 | -17.3 | -19.7 | no *mir-795* | no *mir-x* |
| *C. briggsae* | ACACCCCUUUUAUACAACCAUUCUGCCUCU | 491 | **-29.0** | -16.9 | -17.3 | -19.7 | no *mir-795* | no *mir-x* |
| *C. remanei* | GACCCCUUUUUAUACAACCAUUCUGCCUCA | 817 | **-29.9** | -17.8 | -20.3 | -17.6 | no *mir-795* | no *mir-x* |
| *C. latens* | GACCCCUUUUUAUACAACCAUUCUGCCUCA | 804 | **-29.9** | -17.8 | -20.3 | -17.6 | no *mir-795* | no *mir-x* |
| *C. wallacei* | ACACCCUUUUUAUACAACCGUUCUGCCUCA | 896 | **-31.0** | -18.4 | -19.9 | -20.0 | no *mir-795* | no *mir-x* |
| *C. tropicalis* | ACCCCCAAUUUAUACAACCGUUCUGCCUCA | 897 | **-31.0** | -18.4 | -19.9 | -20.0 | no *mir-795* | no *mir-x* |
| *C. doughertyi* | CCACCCAAUUUAUACAACCAUUCUGCCUCU | 646 | **-29.0** | -17.5 | -19.0 | -20.6 & -19.5 | no *mir-795* | no *mir-x* |
| *C. brenneri* | ACCCCCUUUUUAUACAACCAUUCUGCCUCU | 916 | **-29.0** | -16.9 | -21.5 | -17.8 | no *mir-795* | no *mir-x* |
| *C. inopinata* | GCAAACCUCUUAUACAACCGUUCUGCCUCG | 851 | **-30.5** | -19.9 | -17.8 | -19.6 | no *mir-795* | no *mir-x* |
| *C. elegans* | AAUCCCUUUUUAUACAACCAUUCUGCCUCU | 757 | **-29.0** | -16.9 | -20.4 | -16.9 | -15.8 | no *mir-x* |
| *C. kamaaina* | GCACCCUUUUUAUACAACCAUUCUGCCUCC | 764 | **-28.6** | -17.9 | -20.3 | -16.5 | no *mir-795* | no *mir-x* |
| *C. waitukubuli* | CCCAACUUUAUAUACAAUCAUUCUGCCUCA | 775 | No *let-7* | -19.7 | **-22.1** | -18.4 | no *mir-795* | -21.4 |
| *C. panamensis* | UUUGUUGACAAAUGAGUCCCGACUGCCUCA | 1736 | No *let-7* | -19.9, -22.6, -25.0, **-25.8** & -23.6 | -25.2 | -20.8 | no *mir-795* | -22.5 |
| *C. nouraguensis* | CCCACAACUUUAUACAACCAUUCUGCCUCA | 719 | No *let-7* | -21.3 | -21.6 & -23.7 | -17.8 | no *mir-795* | -23.3, **-23.8** & -20.4 |
| *C. becei* | CCCACAACUUUAUACAACCACUCUGCCUCA | 729 | No *let-7* | **-21.3** | -20.8 | -20.4 | no *mir-795* | -18.8 & -22.1 |
| *C. macrosperma* | CCCCCAACUUUAUACAACCGUUCUGCCUCA | 743 | **-31.0** | -19.6 | -21.8 | -20.0 | no *mir-795* | -21.1 |
| *C. sulstoni* | GCUCGUUUUUAAAUAUUAUACUCUGCCUCA | 655 | No *let-7* | -21.0 | **-24.6** | -23.1 | no *mir-795* | -21.7 |
| *C. afra* | CUGUGCAACCCCUUAUUAUACUCUGCCUCA | 699 | No *let-7* | -19.6 | **-23.2** | -19.6 | no *mir-795* | -17.9 |
| *C. japonica* | GUACCAUUUUAUACCCCCAGUUCUGCCUCA | 654 | No *let-7* | -22.7 | -18.9 | -18.6 | no *mir-795* | **-24.2** |
| *C. uteleia* | CCCAAUAUUUUAUACAACCGUUCUGCCUCG | 832 | **-30.6** | no *mir-48* | -22.2 | no *mir-241* | no *mir-795* | no *mir-x* |
| *C. guadeloupensis* | UCCGUUUUAUUAUACAACCAUUCUGCCUCU | 651 | **-29.9** | no *mir-48* | -20.7 | no *mir-241* | no *mir-795* | no *mir-x* |
| *C. castelli* | CCACCCAUUUUUUACAACCGUUCUGCCUCU | 461 | **-27.1** | no *mir-48* | -20.5 | no *mir-241* | no *mir-795* | no *mir-x* |
| *C. angaria* | Unable to identify due to sequence gaps | not available | not available | no *mir-48* | not available | no *mir-241* | no *mir-795* | no *mir-x* |
| *C. quiockensis* | ACCACCCUUUUUUACAACCGUUCUGCCUCC | 431 | **-26.7** | no *mir-48* | -20.1 | no *mir-241* | no *mir-795* | no *mir-x* |
| *C. vivipara* | GAACUUUUUUAUACAACCGUACCUGCCUCA | 972 | **-29.8** | no *mir-48* | -20.3 | no *mir-241* | no *mir-795* | no *mir-x* |
| *C. virilis* | AAAAUCCUAGAAUACAACCGUUCUGCCUCG | 676 | **-28.9** | no *mir-48* | -21.0 | no *mir-241* | no *mir-795* | no *mir-x* |
| *C. plicata* | GAAACCCUUUUAUACAACCAUUCUGCCUCA | 742 | **-29.9** | no *mir-48* | -20.0 | no *mir-241* | no *mir-795* | no *mir-x* |
| *C. bovis* | GAACCCAUUGGAUACAACCGUUCUGCCUCG | 1004 | **-29.2** | no *mir-48* | -21.0 | no *mir-241* | no *mir-795* | no *mir-x* |
| *C. parvicauda* #1 | ACCACUCGUAUACAACCACUCUCUGCCUCA | 374 | **-28.3** | no *mir-48* | no *mir-84* | no *mir-241* | no *mir-795* | no *mir-x* |
| *C. parvicauda* #2 | CCACCCUGAUUAUACAACCAUUCUGCCUCU | 408 | **-29.6** | no *mir-48* | no *mir-84* | no *mir-241* | no *mir-795* | no *mir-x* |
| *C. monodelphis* | CAAAUAUGUCUACAACCGCCACCUGCCUCA | 2300 | **-26.8** | no *mir-48* | -19.9 | no *mir-241* | no *mir-795* | no *mir-x* |

**Table S4. *lin-41* GU-SM+SUP sequences and MFEs with all *let-7-family* microRNAs.**

List of all *lin-41* GU-SM+SUP sequences in all *Caenorhabditis* species used in this study, the position of the GU-SM relative to the stop codon, and all predicted MFEs (kcal/mol) of each GU-SM+SUP sequence hybridized with each respective *let-7-family* microRNA. MFEs in bold = lowest energy (most favorable). Outlined in the green box are species recovered in the *Elegans* group. Outlined in the red box are species recovered in the *Japonica* group.

| Strain name | Strain description | Genotype |
|---|---|---|
| VT1367 | WT *C. elegans* expressing *col-19::GFP* | maIs105[col-19::GFP] V |
| DG3913 | *C. elegans* endogenous GFP::LIN-41 tag | *lin-41(tn1541) I* |
| NKZ35 | Inbred *C. inopinata* wild isolate | WT |
| QG122 | *C. kamaaina* wild isolate | WT |
| NIC564 | *C. waitukubuli* wild isolate | WT |
| QG702 | *C. panamensis* wild isolate | WT |
| JU2079 | Inbred *C. nouraguensis* wild isolate | WT |
| QG704 | *C. becei* wild isolate | WT |
| JU2083 | Inbred *C. macrosperma* wild isolate | WT |
| JU2788 | Inbred *C. sulstoni* wild isolate | WT |
| VT3978 | *C. sulstoni* endogenous GFP::LIN-41 tag. Original strain: JU2788 | *lin-41(ma513)* |
| JU1286 | Inbred *C. afra* wild isolate | WT |
| DF5081 | Inbred *C. japonica* wild isolate | WT |

**Table S5. List of *Caenorhabditis* strains used in this study.**

| Oligo ID | Name | Sequence |
|---|---|---|
| oCN480 | crRNA *C. sulstoni dpy-10* | IDT Alt-R™ CRISPR crRNA /AltR1/rGrCrCrUrCrCrGrUrArGrGrCrUrCrCrGrCrGrArGrGrUrUrUrUrArGrArGrCrUrArUrGrCrU/ AltR2/ |
| oCN486 | cs-lin-14 RNAi F | TAAGCAGGTACCCAACAGCAACCAACCCGTTTGTG |
| oCN487 | cs-lin-14 RNAi R | TAAGCAGGTACCTTACTGCGATGGAGGCGGCGAC |
| oCN488 | cs-lin-28 RNAi F | TAAGCAGGTACCGCCAACGCCGCGCTACTTTG |
| oCN489 | cs-lin-28 RNAi R | TAAGCAGGTACCATTCGTCGTCCGCGTCCTTC |
| oCN490 | cs-lin-41-RNAi F | TAAGCAGGTACCGAAACGCTCCCGGACTGGAG |
| oCN491 | cs-lin-41 RNAi R | TAAGCAGGTACCCTAGAAGACACGGATGCAGTTGTTGC |
| oCN492 | cs-hbl-1 RNAi F | TAAGCAGGTACCCGAAGCCTCCATGCATAGGGTC |
| oCN493 | cs-hbl-1 RNAi R | TAAGCAGGTACCTTAGTGGTGTTGCGTTGGGCAG |
| oCN494 | cs-lin-29 RNAi F | TAAGCAGGTACCATCCACATTTGTCCGTTTTGCGG |
| oCN495 | cs-lin-29 RNAi R | TAAGCAGGTACCCTTCATGTGATTGATGATACTTTGCGG |
| oCN501 | cs-unc-22 RNAi F | TAAGCAGGTACCCACCCTCACTGCCACGAATGCG |
| oCN502 | cs-unc-22 RNAi R | TAAGCAGGTACCGTGATCTCCCTTGTTGAGCGAGTCG |
| oCN532 | cs-lin-46 RNAi F | TAAGCAGGTACCATGAATTCTGCCGGAGGTTTCCAC |
| oCN533 | cs-lin-46 RNAi R | TAAGCAGGTACCTCAGAAGGCGGAAAGTGGAAGAG |
| oCN586 | ce-lin-41 RNAi F | TAAGCAAAGCTTGACATGTGTCTGGCGTTGAAAATG |
| oCN587 | ce-lin-41 RNAi R | TAAGCAGGTACCGAAGACACGGATGCAATTGTTTCCG |
| oCN545 | 5' *C. sulstoni lin-41* HR+GFP | CTTGGAATCACTTGTTTGTGTCCCGAAACAAAGGAAAAATGAGTAAAGGAGAAGAACTTT |
| oCN546 | GFP+linker | TCCACCTCCAGATCCACCTCCAGATCCACCTCCAGATTTGTATAGTTCGTCCATGCCATG |
| oCN547 | 3' *C. sulstoni lin-41* linker+HR | TCAAGAGTCGGGCATCCGTTCGACACGTTAGACGCAGATCCACCTCCAGATCCACCTCCA |
| oCN551 | crRNA *C. sulstoni lin-41* N-term | IDT Alt-R™ CRISPR crRNA /AltR1/rUrGrUrCrCrCrGrArArArCrArArArGrGrArArArArGrUrUrUrUrArGrArGrCrUrArUrGrCrU/ AltR2/ |
| oCN599 | cm-let-7 geno F | GTACTGCCACTCCAAGAGAGTCG |
| oCN600 | cm-let-7 geno R | ATTGCTCAAAGAGCCGGAAAACTG |
| | oligo (dT) 20 | TTTTTTTTTTTTTTTTTTTT |
| | tracrRNA | IDT Alt-R™ CRISPR tracrRNA |

**Table S6. List of oligos used in this study.**

| **Allele name** | **crRNA** | **Mutation** |
|---|---|---|
| ma513 | cr39 | CSP32.scaffold00176:<br>35802<ccgaaacaaaggaaa**[aatgagtaaaggagaagaacttttcactggagttgtcccaattcttgttgaattagatggtgatgttaatgggcacaaattttctgtcagtggagagggtgaaggtgatgcaacatacggaaaacttacccttaaatttatttgcactactggaaaactacctgttccatgggtaagtttaaacatatatatactaactaaccctgattatttaaattttcagccaacacttgtcactactttctgttatggtgttcaatgcttctcgagatacccagatcatatgaaacggcatgactttttcaagagtgccatgcccgaaggttatgtacaggaaagaactatatttttcaaagatgacgggaactacaagacacgtaagtttaaacagttcggtactaactaaccatacatatttaaattttcaggtgctgaagtcaagtttgaaggtgataccctgttaatagaatcgagttaaaaggtattgatttaaagaagatggaaacattcttggacacaaattggaatacaactataactcacacaatgtatacatcatggcagacaaacaaaagaatggaatcaaagttgtaagtttaaacatgattttactaactaactaatctgatttaaattttcagaacttcaaaattagacacaacattgaagatggaagcgttcaactagcagaccattatcaacaaaatactccaattggcgatggccctgtccttttaccagacaaccattacctgtccacacaatctgccctttcgaaagatcccaacgaaaagagagaccacatggtccttcttgagtttgtaacagctgctgggattacacatggcatggacgaactatacaaatctggaggtggatctggaggtggatctggaggtggatct]**gcgtctaacgtgtcg>35834 |

nucleotide position # <flanking genomic sequence [**inserted sequence**] flanking genomic sequence> nucleotide position #

**Table S7. *C. Sulstoni GFP::lin-41* allele generated by CRISPR/Cas9 editing.**

### *C. elegans*

| Hours after plating (20ºC) | *let-7* | *mir-48* | *mir-241* | *mir-84* | *mir-795* | *lin-4* |
|---|---|---|---|---|---|---|
| **4** | 8.26 | 13.92 | 0.82 | 607.88 | 27.46 | 36.91 |
| **8** | 26.19 | 62.05 | 5.40 | 995.28 | 71.96 | 1325.91 |
| **12** | 35.89 | 521.53 | 45.82 | 2031.43 | 107.41 | 12934.33 |
| **16** | 23.31 | 506.03 | 56.24 | 2470.66 | 114.95 | 13603.64 |
| **20** | 59.43 | 1211.79 | 105.90 | 3125.01 | 135.16 | 15425.77 |
| **24** | 237.51 | 3014.80 | 220.94 | 5235.47 | 155.93 | 18351.98 |
| **28** | 9225.93 | 7403.50 | 470.66 | 9463.55 | 153.15 | 22017.58 |
| **32** | 11017.43 | 8337.78 | 488.08 | 10833.18 | 213.00 | 18990.07 |
| **36** | 13410.50 | 8433.07 | 546.45 | 9453.36 | 121.06 | 17490.85 |
| **40** | 12558.71 | 7953.44 | 434.62 | 7327.59 | 118.34 | 18213.08 |
| **44** | 13562.47 | 7563.44 | 384.41 | 7617.09 | 114.97 | 15473.48 |
| **48** | 13595.46 | 8027.35 | 348.12 | 8205.65 | 108.20 | 11886.52 |

### *C. macrosperma*

| Hours after plating (25ºC) | *let-7* | *mir-48* | *mir-241* | *mir-84* | *mir-x* | *lin-4* |
|---|---|---|---|---|---|---|
| **8** | 7.57 | 9.85 | 117.4 | 43.84 | 84.34 | 8275.34 |
| **17** | 6.94 | 49.48 | 837.47 | 367.21 | 794.31 | 15548.37 |
| **22** | 26.25 | 127.79 | 2120.78 | 1072.58 | 2454.5 | 19495.40 |
| **29** | 71.5 | 213.14 | 2531.63 | 1972.41 | 4231.99 | 18170.43 |
| **33** | 48.82 | 150.54 | 1441.63 | 1644.94 | 3607.63 | 13324.01 |

### *C. sulstoni*

| Hours after plating (25ºC) | *mir-48* | *mir-241* | *mir-84* | *mir-x* | *lin-4* |
|---|---|---|---|---|---|
| **4** | 10.39 | 11.07 | 110.33 | 43.98 | 40.24 |
| **8** | 248.11 | 303.02 | 1049.45 | 347.49 | 11992.21 |
| **12** | 574.93 | 339.23 | 1795.61 | 664.38 | 10586.87 |
| **16** | 4260.99 | 1306.80 | 4720.21 | 2105.92 | 12745.80 |
| **20** | 6529.49 | 1726.07 | 5079.74 | 2517.81 | 12259.29 |
| **24** | 8105.97 | 1293.41 | 5160.28 | 2623.18 | 11038.84 |
| **28** | 7545.82 | 1294.12 | 5319.99 | 2754.10 | 7493.32 |
| **32** | 5131.69 | 787.44 | 3826.11 | 1678.41 | 3708.38 |

**Table S8. Normalized (RPM) small RNA sequencing expression numbers used for Figures 3 and S4.**

| temperature | *C. elegans* Hours after plating L1 molt | L2 molt | L3 molt | L4 molt | *C. macrosperma* Hours after plating L1 molt | L2 molt | L3 molt | L4 molt | *C. sulstoni* Hours after plating L1 molt | L2 molt | L3 molt | L4 molt |
|---|---|---|---|---|---|---|---|---|---|---|---|---|
| **15ºC** | 20.5 | 32 | 44 | 59 | n/d | n/d | n/d | n/d | 50 | n/d | n/d | 144 |
| **20ºC** | 16 | 28 | 40 | 50 | n/d | n/d | n/d | n/d | 16 | 26 | 34 | 48 |
| **25ºC** | 12.5 | 20 | 31 | 41 | 16 | 20 | 24 | 32 | 10 | 16 | 22 | 28 |
| **30ºC** | 0 | 0 | 0 | 0 | n/d | n/d | n/d | n/d | 10 | 14 | 17 | 23 |
| **33ºC** | 0 | 0 | 0 | 0 | n/d | n/d | n/d | n/d | 10 | 14 | 17 | 23 |
| **35ºC** | 0 | 0 | 0 | 0 | n/d | n/d | n/d | n/d | 9 | 0 | 0 | 0 |

n/d = not determined

**Table S9. Timing numbers of developmental growth rates use for Figures 3, S3, and S4.**

**C. elegans: alae completeness**

| | | No alae | Gapped alae | Complete alae |
|---|---|---|---|---|
| # L4s | *lin-41* RNAi | 5 | 15 | 0 |
| | control | 12 | 0 | 0 |
| # adults | *lin-41* RNAi | 0 | 19 | 4 |
| | control | 0 | 1 | 10 |

**C. sulstoni: alae completeness**

| | | No alae | Gapped alae | Complete alae |
|---|---|---|---|---|
| # L4s | *lin-14* RNAi | 0 | 11 | 4 |
| | *lin-28* RNAi | 0 | 5 | 6 |
| | *lin-41* RNAi | 34 | 8 | 8 |
| | *hbl-1* RNAi | 0 | 16 | 1 |
| | control | 47 | 1 | 0 |
| # adults | *lin-29* RNAi | 0 | 17 | 0 |
| | *lin-41* RNAi | 33 | 10 | 0 |
| | *lin-46* RNAi | 0 | 9 | 38 |
| | control | 0 | 7 | 57 |

**C. sulstoni: # branches**

| | | 0 | 1 | 2 | 3 | 4 |
|---|---|---|---|---|---|---|
| # adults | *lin-46 RNAi* | 21 | 11 | 8 | 3 | 4 |
| | control | 39 | 19 | 5 | 1 | 0 |

**Table S10. RNAi quantification numbers used for Figures 4 and 7.**

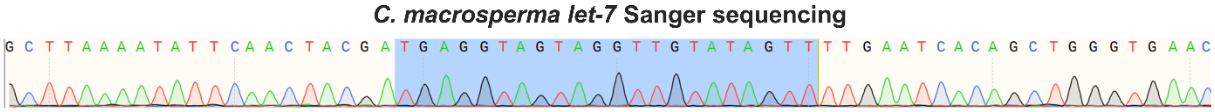

**Figure S1. Sanger sequencing of *let-7* microRNA sequence in *C. macrosperma*.**
Sanger sequencing confirming the *let-7* microRNA sequence (highlighted in blue) in the *C. macrosperma* genome.

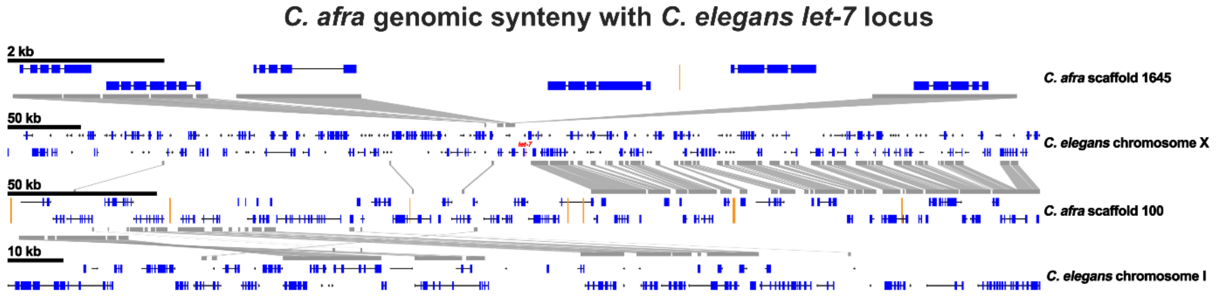

**Figure S2. The genomic region containing the *let-7* sequence in *C. elegans* is syntenic to multiple scaffolds in *C. afra*.**

A portion of *C. elegans* chromosome X that is 3' of the *let-7* sequence (left side of *let-7* in the image) is syntenic to *C. afra* scaffold 1645. A portion of *C. elegans* chromosome X that is 5' of the *let-7* sequence (right side of *let-7* in the image) is syntenic to a portion of *C. afra* scaffold 100. Another portion of *C. afra* scaffold 100 is also syntenic to a portion of *C. elegans* chromosome I. Regions with sequence similarity are outlined in gray. Sequence gaps are shown in orange. *let-7* is shown in red (center of the *C. elegans* chromosome X portion shown).

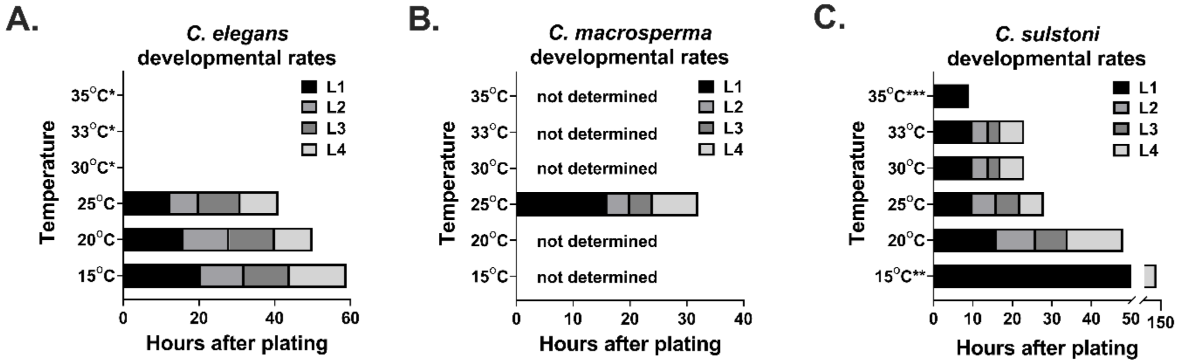

**Figure S3. Developmental growth rates in *C. elegans*, *C. macrosperma*, and *C. sulstoni.***
Developmental growth rates of L1 arrested *C. elegans* (A), *C. sulstoni* (B), and *C. macrosperma* (C) at various temperatures. *animals did not exit L1 arrest. **animal development was asynchronous; timing of first animals reaching adulthood is shown. ***animals failed to exit L1 molt.

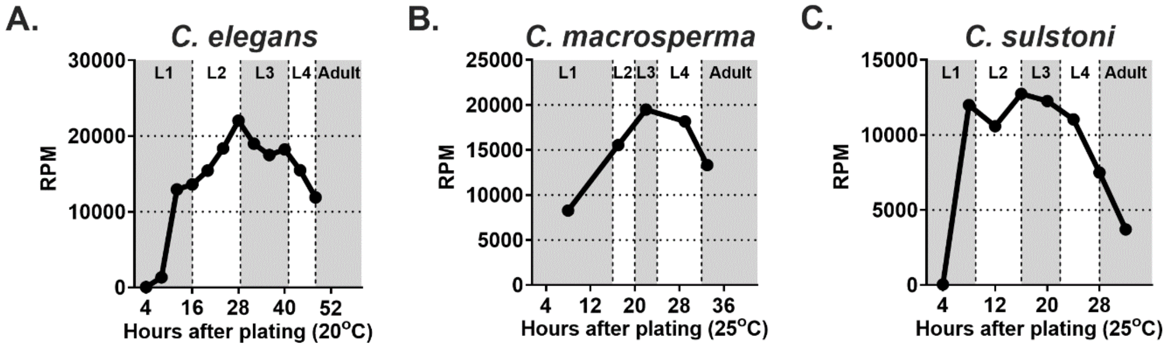

**Figure S4. The temporal expression patterns of *lin-4* microRNA in *C. elegans*, *C. macrosperma*, and *C. sulstoni.***

Small RNA sequencing data showing expression of *lin-4* microRNA throughout *C. elegans* (A), *C. macrosperma* (B), and *C. sulstoni* (C) development. RPM refers to reads per million.

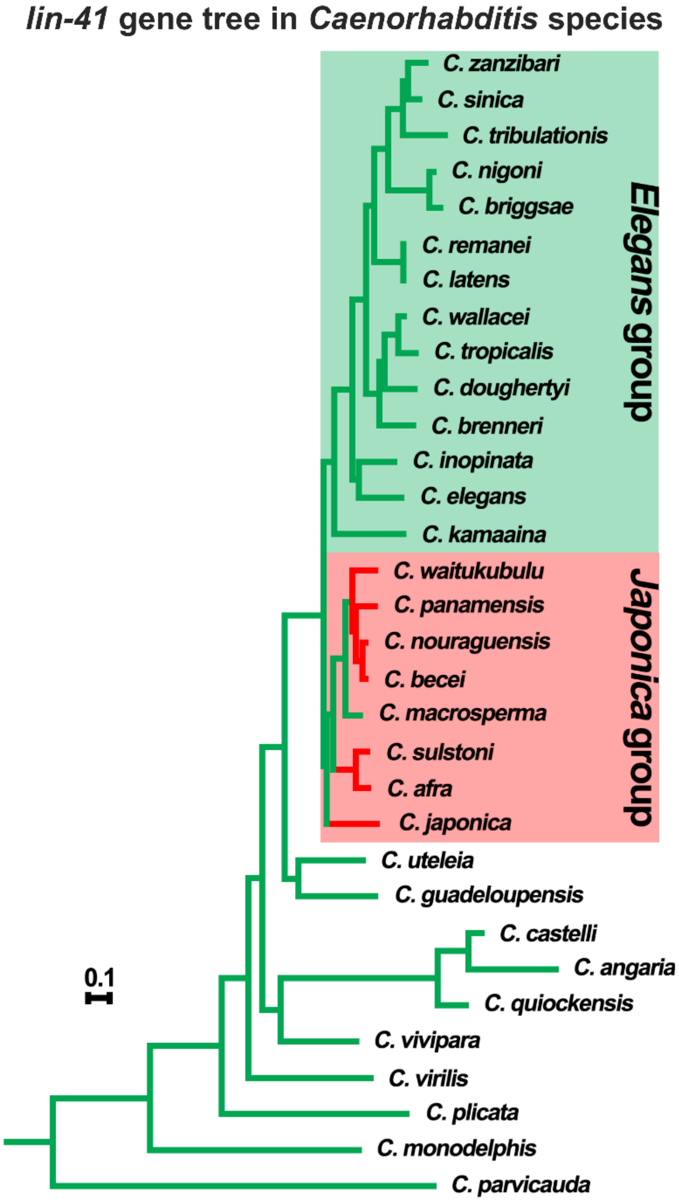

**Figure S5. *lin-41* phylogeny in *Caenorhabditis.***

Gene tree of *lin-41* protein sequence in *Caenorhabditis* species based on orthology clustering provided by the *Caenorhabditis* Genomes Project (Stevens, 2020). Green branches highlight species that are either predicted or confirmed to have *let-7*. Red branches highlight species that are either predicted or confirmed to lack *let-7*. Outlined in the green box are species belonging to the *Elegans* group. Outlined in the red box are species belonging to the *Japonica* group. Branch lengths are in substitutions per site. Scale bar is shown.

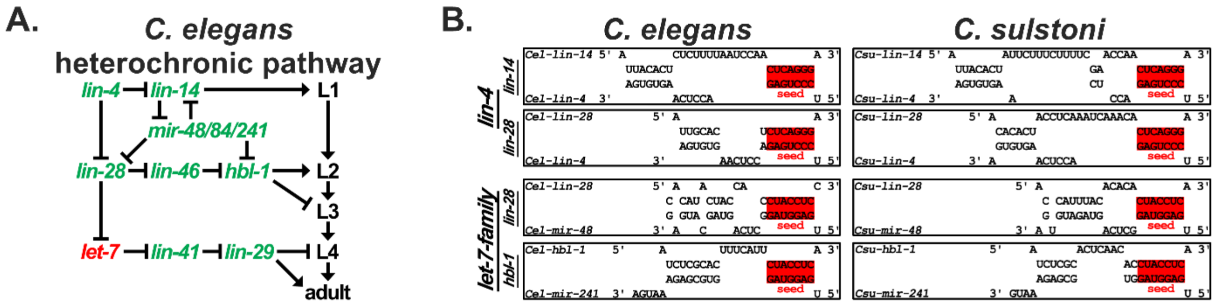

**Figure S6. Integration of *lin-4* and the *let-7-family* microRNAs into the heterochronic pathway is conserved between *C. elegans* and *C. sulstoni.***

(A) Genetic diagram of the heterochronic pathway in *C. elegans*. Genes in green font are conserved in *C. sulstoni*. Gene in red font is not found in *C. sulstoni*. Adapted from Resnick et al., 2010.

(B) Predicted base pairing of *lin-4* with a representative complementary site in the *lin-14* 3' UTR and with the complementary site in the *lin-28* 3' UTR (top two rows), predicted base pairing of *mir-48* with the complementary site in the *lin-28* 3' UTR (third row), and predicted base pairing of *mir-241* with a representative complementary site in the *hbl-1* 3' UTR (fourth row) in *C. elegans* (left column) and *C. sulstoni* (right column).

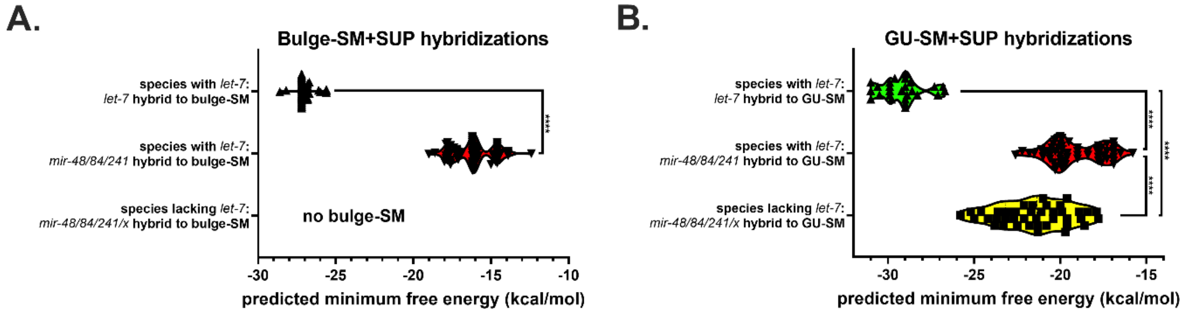

**Figure S7. Among *let-7-family* microRNAs, *let-7* has the most favorable hybridization to the *lin-41* SMs.**

Predicted hybridization MFEs of *let-7-family* microRNAs with the *lin-41* bulge-SM+SUP (A) and GU-SM+SUP (B) in *Caenorhabditis* species. Statistical significance was determined using a two-tailed Student's t-test. ***** $P \leq 0.0001$.

**WORKS CITED**

**De Wit, E., Linsen, S. E., Cuppen, E. and Berezikov, E.** (2009). Repertoire and evolution of miRNA genes in four divergent nematode species. *Genome Res* **19,** 2064-74.

**Lau, N. C., Lim, L. P., Weinstein, E. G. and Bartel, D. P.** (2001). An abundant class of tiny RNAs with probable regulatory roles in Caenorhabditis elegans. *Science* **294,** 858-62.

**Lim, L. P., Lau, N. C., Weinstein, E. G., Abdelhakim, A., Yekta, S., Rhoades, M. W., Burge, C. B. and Bartel, D. P.** (2003). The microRNAs of Caenorhabditis elegans. *Genes Dev* **17,** 991-1008.

**Moss, E. G.** (2007). Heterochronic genes and the nature of developmental time. *Curr Biol* **17,** R425-34.

**Pasquinelli, A. E., Reinhart, B. J., Slack, F., Martindale, M. Q., Kuroda, M. I., Maller, B., Hayward, D. C., Ball, E. E., Degnan, B., Muller, P., et al.** (2000). Conservation of the sequence and temporal expression of let-7 heterochronic regulatory RNA. *Nature* **408,** 86-9.

**Reinhart, B. J., Slack, F. J., Basson, M., Pasquinelli, A. E., Bettinger, J. C., Rougvie, A. E., Horvitz, H. R. and Ruvkun, G.** (2000). The 21-nucleotide let-7 RNA regulates developmental timing in Caenorhabditis elegans. *Nature* **403,** 901-6.

**Resnick, T. D., Mcculloch, K. A. and Rougvie, A. E.** (2010). miRNAs give worms the time of their lives: small RNAs and temporal control in Caenorhabditis elegans. *Dev Dyn* **239,** 1477-89.

**Rougvie, A. E.** (2001). Control of developmental timing in animals. *Nat Rev Genet* **2,** 690-701.

**Ruby, J. G., Jan, C., Player, C., Axtell, M. J., Lee, W., Nusbaum, C., Ge, H. and Bartel, D. P.** (2006). Large-scale sequencing reveals 21U-RNAs and additional microRNAs and endogenous siRNAs in C. elegans. *Cell* **127,** 1193-207.

**Shi, Z., Montgomery, T. A., Qi, Y. and Ruvkun, G.** (2013). High-throughput sequencing reveals extraordinary fluidity of miRNA, piRNA, and siRNA pathways in nematodes. *Genome Res* **23,** 497-508.

**Stevens, L.** (2020). CGP orthology clustering. *Zenodo*. https://doi.org/10.5281/zenodo.4068211
